# Molecular Mechanisms of DNMT3A-3L-Mediated *de novo* DNA Methylation on Chromatin

**DOI:** 10.1101/2025.06.10.658647

**Authors:** Yan Yan, X. Edward Zhou, Stacey L Thomas, Minmin Liu, Gan-Qiang Lai, Evan J Worden, Peter A Jones, Ting-Hai Xu

**Affiliations:** Edward A. Doisy Department of Biochemistry and Molecular Biology, Saint Louis University School of Medicine, St. Louis, MO, 63104, USA; Department of Structural Biology, Van Andel Institute, Grand Rapids, MI 49503, USA; Department of Epigenetics, Van Andel Institute, Grand Rapids, MI 49503, USA

## Abstract

*De novo* DNA methylation is a fundamental epigenetic process essential for early development. It is mediated by DNA methyltransferase DNMT3A and DNMT3B, in cooperation with catalytically inactive DNMT3L. Abnormal DNA methylation contributes to genomic instability and oncogenesis, yet the mechanisms governing its establishment and regulation remain unclear. Here, we present high-resolution cryo-EM structures of the nucleosome-bound full-length DNMT3A2-3L and its oligomeric assemblies in the nucleosome-free state. Strikingly, DNMT3L C-terminal “Switching Helix” displayed a distinct conformation with a 180° rotational difference compared to its DNMT3B3 counterpart in the presence of nucleosome, preventing direct interactions with the nucleosome acidic patch. Instead, DNMT3L ADD domain promotes nucleosome binding, while DNMT3A PWWP domain inhibits it, indicating multi-layer regulation. Additionally, the novel oligomeric arrangement of DNMT3A2-3L in nucleosome-free states highlights its dynamic assembly and potential allosteric regulation. Furthermore, we identified the critical role of DNMT3L as a sensor of histone modifications, guiding the complex’s chromatin interaction. These findings uncover a previously unknown mechanism by which DNMT3A-3L mediates *de novo* DNA methylation on chromatin through complex reorganization, histone tail sensing, dynamic DNA search, and nucleosome engagement, providing key insights into epigenetic regulation.

## Introduction

The epigenome is a sophisticated regulatory network involving chemical modifications of DNA and histones that are crucial for controlling chromatin architecture and gene expression. Key mechanisms include DNA methylation, histone modification, and chromatin remodeling. Among these, DNA methylation, which predominantly occurs on the 5-carbon of cytosine within CpG dinucleotides, is essential for normal development and differentiation. It influences biological processes such as transcriptional silencing, genomic imprinting, and X-chromosome inactivation^1^. DNA methylation patterns are largely erased and re-established between generations^2^. These patterns are orchestrated by two *de novo* methyltransferases, DNMT3A and DNMT3B, which share significant sequence and structural similarities^1^, and are assisted by catalytically inactive accessory paralogues, such as DNMT3L and DNMT3B3. Notably, DNMT3L is primarily expressed in embryonic stem (ES) cells^3^, where DNA methylation patterns are first established. Interestingly, DNMT3A2 and DNMT3L are repressed during differentiation (**Fig. 1a**), leaving DNMT3B3 as a functional substitution of DNMT3L in somatic cells^4,5^, This distinct cell type-specific expression pattern highlights the specialized role of DNMT3A2 and DNMT3L in mediating *de novo* DNA methylation in early embryonic development. This raises the question of how *de novo* DNA methylation is established and regulated in the early stages of embryogenesis, and how DNMT3L contributes to this process in a unique way. Although partial structures of isolated DNMT3 domains have provided insights into their DNA binding and catalysis^6–9^, the precise molecular mechanism underlying *de novo* DNA methylation within the chromatin context, particularly the DNMT3L-dependent regulation, remains unclear. Our previous studies have shed light on the aspects of DNMT3B3-mediated DNA methylation^10^, but the specific mechanisms by which DNMT3L-mediated DNA methylation are yet to be defined.

**Fig. 1.**
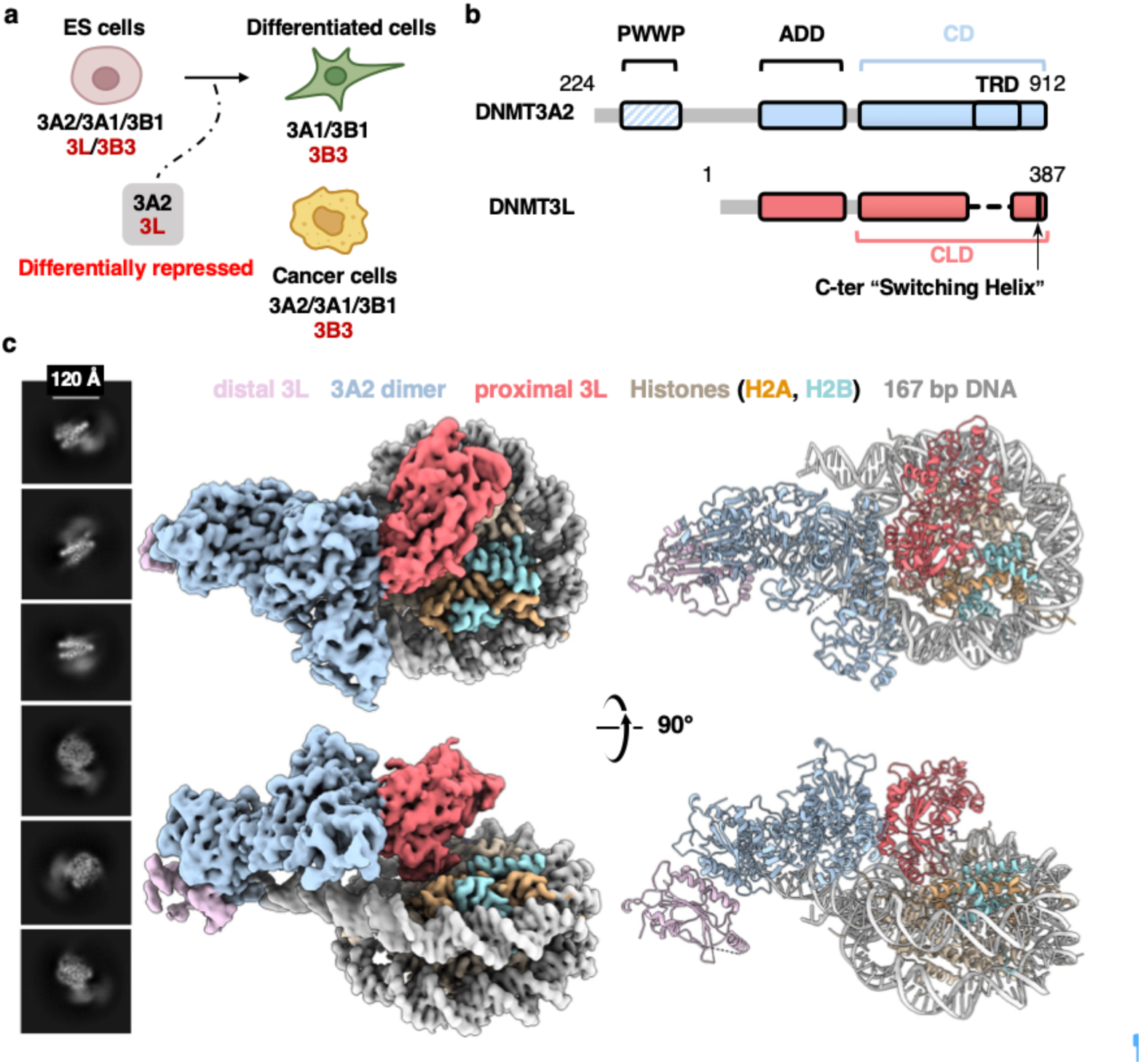
Cryo-EM structure of DNMT3A2-3L in nucleosome-bound state. **a,** The catalytically active *de novo* CpG methyltransferases DNMT3s (3A1, 3A2, and 3B1) and the catalytically inactive accessory DNMT3s (3L and 3B3) are expressed in embryonic stem (ES) cells. DNMT3A2 and DNMT3L are repressed during differentiation. DNMT3B3 is the predominant accessory protein in differentiated cells, while DNMT3A2 is notably overexpressed in cancer. **b,** Color-coded domain architectures of human DNMT3A2-3L complex. **c,** Left, representative 2D class-averages. Right, cryo-EM density map and atomic model of the nucleosome-bound DNMT3A2-3L complex. DNMT3A2-3L proteins were colored according to the domain architecture in panel **b**. Histone H2A and H2B were highlighted with orange and cyan, respectively. The distal DNMT3L was in pink.

DNMT3A2, the predominant DNMT3A isoform in ES cells^11^ and cancer cells^10^ (**Fig. 1a**), contains several functional domains critical for its activity: an N-terminal Pro-Trp-Trp-Pro (PWWP) domain, which recognizes and binds to lysine 36 di-/tri-methylated histone H3 (H3K36me2/3) and DNA^12–16^; a central ATRX-DNMT3-DNMT3L (ADD) domain, which specifically interacts with histone H3 tails when lysine 4 is unmethylated (H3K4me0)^17,18^; and a C-terminal catalytic domain (CD) that harbors conserved motifs responsible for catalyzing the transfer of a methyl group from S-adenosylmethionine (SAM) to the 5-carbon of cytosines in DNA^19^. In contrast, DNMT3L shares sequence homology with DNMT3A but lacks the N-terminal PWWP domain and the target recognition domain (TRD) in the C-terminal catalytic-like domain (CLD) (**Fig. 1b**), and therefore has no DNA methyltransferase activity. However, DNMT3L significantly stimulates DNA methylation by directly interacting with DNMT3A and DNMT3B^5,20^.

In this study, we investigated the regulatory mechanism of DNMT3A2-3L-mediated *de novo* DNA methylation in the context of intact nucleosomes. By assembling a nucleosome core particle (NCP) with the full-length DNMT3A2-3L complex, we determined its cryo-EM structure at a global resolution of 3.10 Å, with sub-regions resolved to 3.08 Å for NCPs and 3.60 Å for DNMT3A2-3L (**Fig. 1c** and **Extended Data Figs. 1-2**). We also determined the cryo-EM structures of the DNMT3A2-3L complex in its nucleosome-free states at resolutions of 3.66 Å for the dodecamer and 3.62 Å for the hexamer (**Fig. 2a-b** and **Extended Data Fig. 3**). These structures revealed a previously uncharacterized oligomeric organization of the DNMT3A2-3L complex and a distinct nucleosome-binding mechanism that is substantially different from that of the DNMT3A2-3B3 complex^10^. Our structural findings, together with supporting biochemical data, suggest a searching model in which DNMT3A2-3L reorganizes its organization, senses histone tails, and dynamically searches for DNA and engages with nucleosomes, providing mechanistic insight into DNMT3A2-3L-mediated *de novo* DNA methylation on chromatin.

**Fig. 2.**
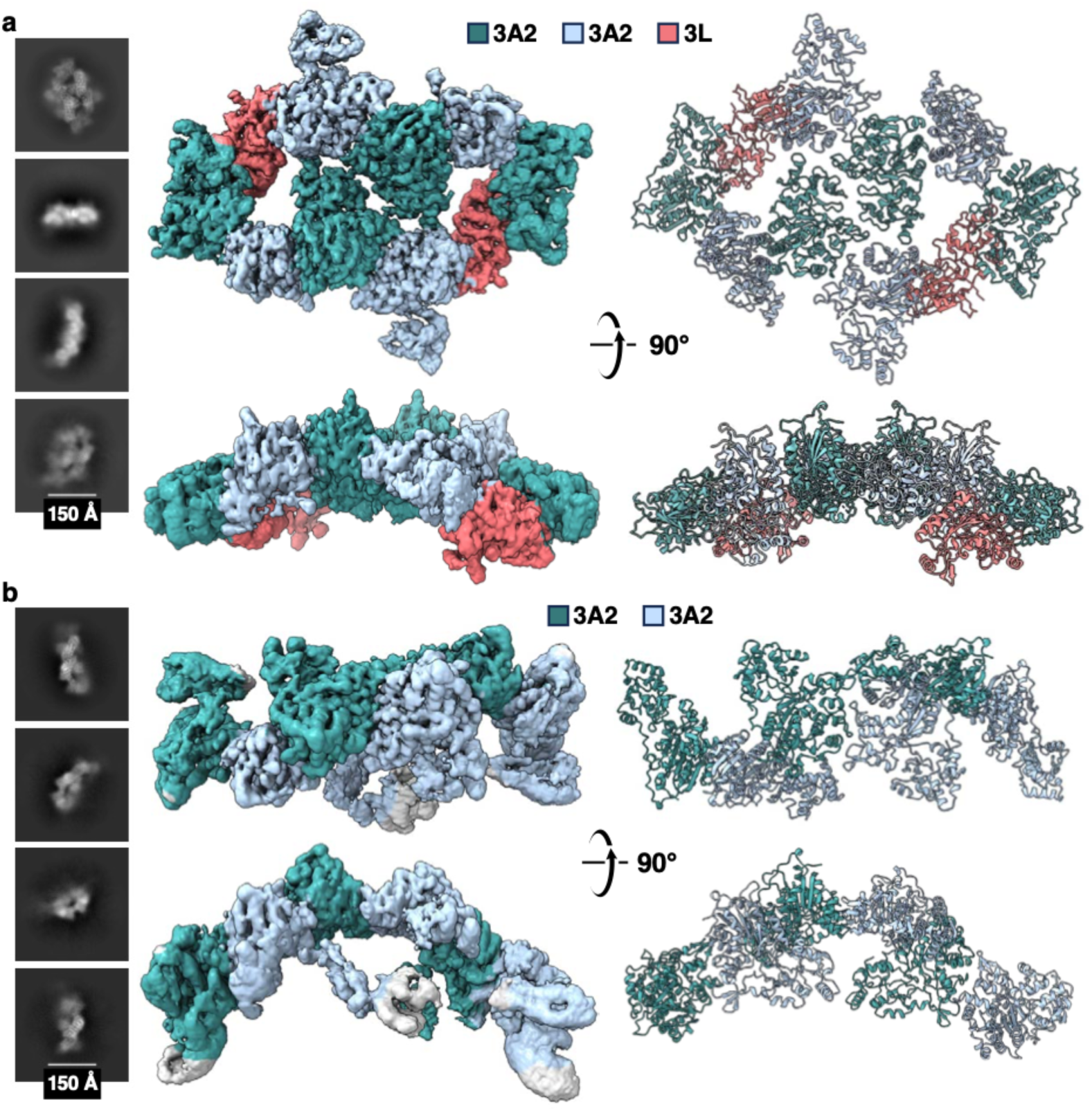
Cryo-EM structures of DNMT3A2-3L in nucleosome-free state. **a, b,** Cryo-EM analysis of the nucleosome-free DNMT3A2-3L complex in dodecamer state (**a**) and in hexamer state (**b**). Left, representative 2D class-averages; right, cryo-EM density maps and the atomic models. DNMT3L were shown in red and DNMT3A2 in sea green and light blue.

### Cryo-EM structures of DNMT3A2–DNMT3L reveal nucleosome binding and oligomeric organization

To reveal the molecular mechanism of DNMT3A-3L binding to nucleosomes, we reconstituted NCPs containing the 147-base-pair (bp) ‘Widom 601’ sequence^21^ and 10 bp linker on each side as previously described (NCP167)^10^, and assembled them with full-length human DNMT3A2-3L complex. Interaction was first confirmed by AlphaScreen luminescence proximity assays (**Extended Data Fig. 1b**). The cryo-EM structure had well-resolved densities for the nucleosome core and the DNMT-bound flanking DNA. To a lesser degree, the structure showed clear density for secondary structure elements of the DNMT3A2 CD domains, the proximal DNMT3A2 ADD domain, and the proximal DNMT3L (**Fig. 1c** and **Extended Data Fig. 4b-c**). The distal DNMT3A2 ADD domain density only demonstrates the overall position (**Extended Data Fig. 4c** inlet), while the PWWP domains and histone tails were unresolved, likely due to the conformational flexibility. Thereby, the cryo-EM structure of this complex shows a clear heterotrimeric DNMT3L-3A2-3A2 arrangement with clear density for the F-F interface helices and the distinct C-terminal helix (we named “Switching Helix”) at the distal interface (**Fig. 1c** and **Extended Data Fig. 4d**). Differences in the C-terminal “Switching Helix” conformation between DNMT3L and DNMT3A, along with the absence of TRD region densities (**Extended Data Fig. 4d**), indicate the presence of a second DNMT3L (distal) to form a more stable and active heterotetramer^8,11,22–24^, consistent with the conformations observed in isolated DNMT3A-3L domain structures^25^. Similar to DNMT3A2-3B3 complex^10^, DNMT3A2-3L also binds to nucleosomes asymmetrically: the proximal DNMT3L engages the nucleosome core, while the distal DNMT3A2 CD interacts with the linker DNA (**Fig. 1c**). DNMT3 complexes are known to adopt two conformations in equilibrium between active and autoinhibitory states. Releasing the autoinhibition state requires the ADD domain to bind to an unmodified histone H3 N terminus^25^. Despite the presence of unmodified H3 tails in our NCPs, the DNMT3A2 ADD domains remained in an autoinhibitory conformation in our cryo-EM structure (**Fig. 1c** and **Extended Data Fig. 4e**). Interestingly, the proximal DNMT3L ADD domain adopted an active conformation (**Extended Data Fig. 4f**), suggesting that engagement with the nucleosome core selectively modulates DNMT3L ADD domain conformations to regulate its binding and activity.

To further gain insights into the DNMT3A assembly, we determined the nucleosome-free structures of DNMT3A2-3L in two oligomeric states. The heterotetramer of the DNMT3A2-3L CD/CLD domains revealed two distinct interaction interfaces: a “flat”, non-polar, heterodimeric interface (3A-3L F-F interface) and a “V-shaped” polar, homodimeric interface (3A-3A R-D interface)^8,9^ (**Extended Data Fig. 4a**). These structural features facilitated the identification of four R-D interfaces in the dodecamer (**Fig. 2a**) and three in the hexamer (**Fig. 2b**), corresponding to four and three DNMT3A2 dimers, respectively. In the dodecamer, regions beyond DNMT3L-3A2-3A2-3A2-3A2 could not be confidently assigned due to resolution limits (**Extended Data Fig. 3e**), and therefore they were omitted from the final model. Since DNMT3L can only form the F-F interface and prevent higher-order oligomerization of DNMT3A2^26^, we hypothesized that the distal regions correspond to two additional DNMT3L molecules, resulting in the DNMT3A2-3L dodecamer complex. In the hexamer, regions beyond the three DNMT3A2 dimers were too weak to confidently assign (**Extended Data Fig. 3f**), and they were omitted from the final model. While DNMT3B homo-oligomerization has been structurally characterized^27^, our cryo-EM analysis provides the first evidence of DNMT3A2-3L forming hetero higher-order oligomers, which may increase the local concentration of DNMT3A2-3L to promote efficient chromatin scanning. Importantly,. this unique novel arrangement enables dynamic assembly and allosteric regulation of the DNMT3A2-3L complex.

These findings indicate that nucleosomes function not only as substrates for the DNMT3A complex but also as regulators for DNMT3A assembly. When bound to nucleosomes, the heterotetrameric arrangement likely stabilizes DNMT3A2-3L while preventing higher-order oligomerization and enabling the complex to adopt a more stable and active conformation^8,11,22–24^. Despite this stabilization, we still did not observe the flipped DNA base in the DNA-engaged DNMT3A2 catalytic center (**Extended Data Fig. 4g**), indicating that nucleosomes binding does not require CpG motifs and is not sequence-specific. This sequence-independent binding may provide a reasonable explanation for the absence of strict sequence patterns in genome-wide DNA methylation.

### DNMT3L C-terminal “Switching Helix” prevents nucleosome acidic patch interaction

Structural comparison of nucleosome-bound DNMT3A2-3L and nucleosome-bound DNMT3A2-3B3 reveals a striking ∼180° rotational difference in the conformation of the DNMT3L C-terminal “Switching Helix” relative to its DNMT3B3 counterpart. This rotated conformation prevents the interactions between the accessory protein DNMT3L and the nucleosome acidic patch (**Fig. 3a**), a key binding site formed by histones H2A and H2B for various nucleosome-interacting proteins, including DNMT3A2-3B3^10^. Cryo-EM density maps further confirmed that, in this rotated conformation, DNMT3L CLD does not directly contact the acidic patch (**Extended Data Fig. 5a-b**).

**Fig. 3.**
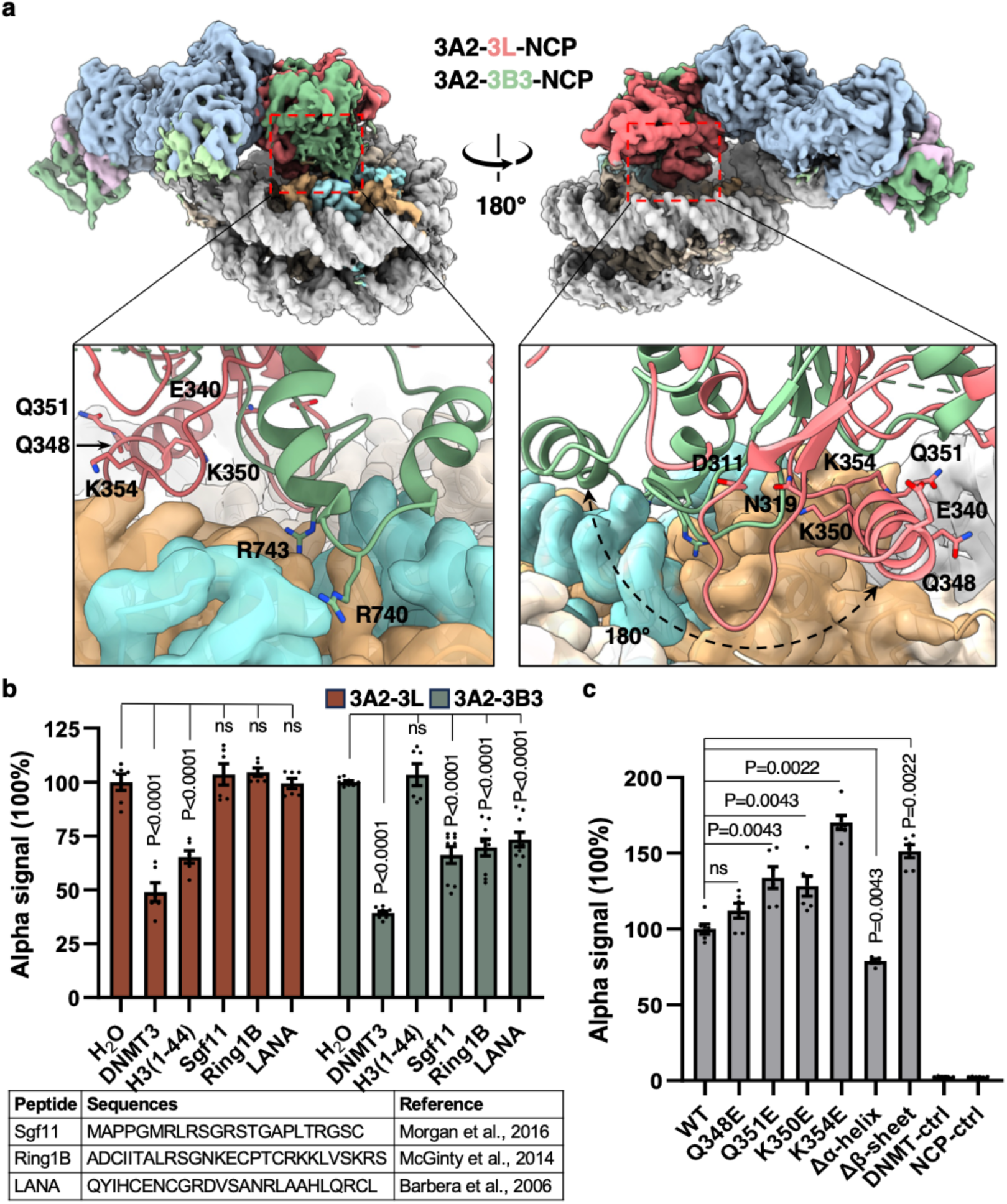
Interactions between DNMT3A2-3L and nucleosomes. **a,** Top, two views (180° apart) of superimposed cryo-EM maps of DNMT3A2-3L-NCP (current) and DNMT3A2-3B3-NCP (EMD-20281). Bottom, close-up views of accessory protein interactions with nucleosome core region (cryo-EM density was shown in **Extended Data Fig. 5a-b**). The nucleosome core particle (NCP) density was shown as transparent. DNMT3L, DNMT3B3, and histone proteins were shown in cartoon models. Color codes were the same as in Fig.1, with DNMT3B3 in green (PDB ID 6PA7). **b,** The acidic patch-interacting peptides and histone H3 peptide competition assay of DNMT3A2-3L (red) and DNMT3A2-3B3 (green). Peptide sequences were listed below. DNMT3, untagged DNMT3A2-3L or DNMT3A2-3B3 complex controls. **c,** AlphaScreen interaction assay between His-tagged wild-type and DNMT3L mutant complexes and biotinylated NCPs. Δα-helix, deletion of residues from E340 to K354. Δβ-sheet, deletion of residues from D311 to N319. Data were presented as mean ± s.e.m., n = 9 (**b** 3A2-3B3 group), and 6 (**b** 3A2-3L group and **c**). ns, not significant. P-values, two-tailed Student’s *t*-test.

To validate this novel observation, we performed nucleosome-binding competition assays using peptides that specifically interact with the acidic patch. Consistent with structural data, acidic patch-interacting peptides reduced the nucleosome binding of the DNMT3A2-3B3 complex, but had no effect on DNMT3A2-3L, suggesting distinct chromatin recruitment mechanisms for the different DNMT3 accessory proteins. Interestingly, the histone H3 peptides selectively competed with DNMT3A2-3L for nucleosome binding (**Fig. 3b**), indicating that histone tails may have a greater role in DNMT3A2-3L recruitment due to its lack of direct interaction with the nucleosome acidic patch. Sequence alignment of DNMT3 family C-terminal regions highlights high conservation between DNMT3A and DNMT3B but not DNMT3L (**Extended Data Fig. 5c**), indicating functional divergence. Closer examination identified residues Q348 and Q351 in DNMT3L as equivalents of the DNMT3B3 “R finger” residues (R740 and R743)^10^. However, due to the unique confirmation of DNMT3L’s C-terminal “Switching Helix”, these residues, along with nearby K350 and K354, do not directly interact with the acidic patch (**Fig. 3a**). Nucleosome-binding assays with DNMT3L-mutant complexes further confirmed that, unlike DNMT3B3, DNMT3L does not directly engage the acidic patch (**Fig. 3c**). Given the proximity of the DNMT3L C-terminal “Switching Helix” and adjacent β-sheets to the nucleosome core (**Fig. 3a**), we evaluated their roles in nucleosome recruitment. Notably, although residues Q348, Q351, K350, and K354 had a subtle impact on nucleosome binding, deletion of the entire C-terminal “Switching Helix” (E340 to K354) - Δα-helix modestly reduced nucleosome binding by approximately 25%, suggesting that this region mediates moderate-range charge and polar interactions. However, these interactions alone appear insufficient to engage nucleosomes, implying the requirement for additional contacts. Conversely, deleting the β-sheet region (D311 to N319)-Δβ-sheet, which contains negatively charged and polar residues (**Extended Data Fig. 5c**), led to a slight increase in nucleosome binding (**Fig. 3c**). Together, these data strongly support the novel finding that DNMT3A2-3L binds to nucleosomes through a mechanism distinct from acidic patch engagement, instead relying on moderate, potentially electrostatic interactions mediated by the C-terminal “Switching Helix” and other nucleosome-proximal regions.

### DNMT3A2-3L binds nucleosome core through the DNMT3L ADD domain

To investigate how DNMT3L contributes to the recruitment of the DNMT3A2–3L complex to nucleosomes, we overlaid the DNMT3A2-3L-nucleosome map with the DNMT3A2-3B3-nucleosome map (EMD-20281). This comparison revealed the additional well-defined density corresponding to the DNMT3L ADD domain, emphasizing its critical role in nucleosome binding (**Fig. 4a**). Structural analysis showed that the DNMT3L ADD domain was anchored by negatively charged residues including E103, which interacts electrostatically with positively charged histone H4 tails particularly K20 (**Fig. 4b**). While eliminating this charge interaction significantly weakened nucleosome binding, the deletion of the DNMT3L ADD domain almost completely abolished it (**Fig. 4c**). This data reinforces the conclusion that DNMT3L ADD domain plays a critical role for nucleosome binding. Surprisingly, deletion of the DNMT3A2 PWWP domain led to a fivefold increase in nucleosome binding (**Fig. 4c**) despite its known affinity for H3K36me2/3 and DNA^12–16^. This observation indicates that the PWWP domain may alter the conformation of DNMT3L N-terminus since DNMT3L is N-terminally His-tagged or may regulate nucleosome interactions by modulating DNA binding dynamics (**Fig. 4d**). The precise regulatory role and mechanism of the PWWP domain await further investigation.

**Fig. 4.**
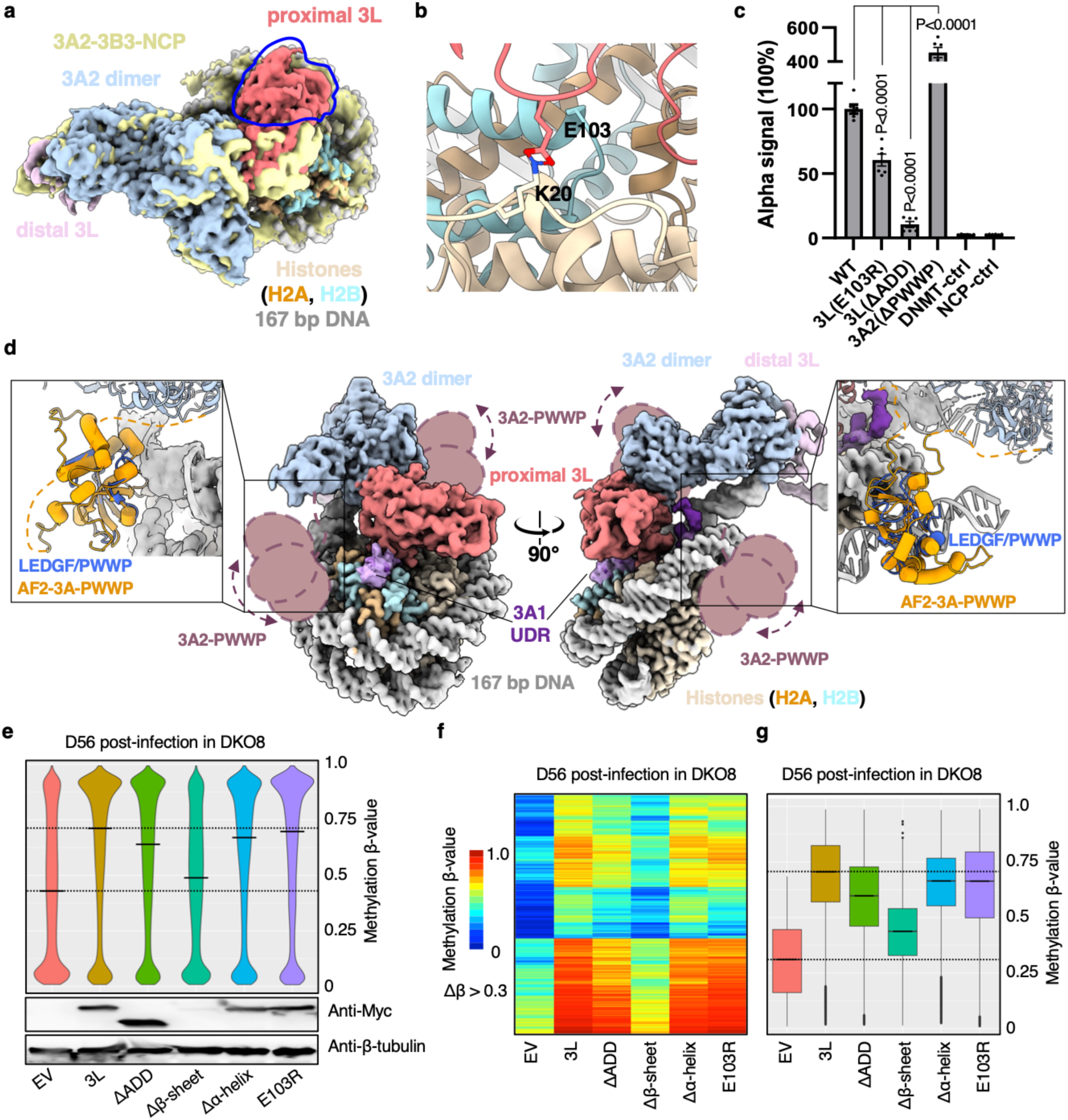
ADD and PWWP domains are important for nucleosome recruitment and regulation. **a,** DNMT3A2-3L-NCP cryo-EM map superimposed on DNMT3A2-3B3-NCP cryo-EM map (EMD-20281). Blue lines highlighted the additional good density corresponding to the DNMT3L ADD domain. **b,** Close-up view of the interaction between DNMT3L ADD loop and histone H4 tails. **c**, AlphaScreen interaction assay between His-tagged wild-type and mutant DNMT3 complexes and biotinylated NCPs. Data were presented as mean ± s.e.m., n = 6. ns, not significant. P-values, two-tailed Student’s *t*-test. **d,** The overall position of the PWWP domain of the distal DNMT3A2 and cryo-EM densities of the DNMT3A1 UDR (EMD-42636, EMD-18778, and EMD-41922) were overlaid with the DNMT3A2-3L-NCP complex. The atomic model and maps were color-coded as in Fig.1, with DNMT3A1 UDRs in varying shades of purple. Inlet, structure of the H3K36me3 modified nucleosome bound to the PWWP domain from LEDGF (PDB ID 6S01 and 8CBN) superimposed with DNMT3A PWWP domain from AF2 model and the nucleosome-bound DNMT3A2-3L complex structure. **e,** Top, violin plots of DNA methylation levels in DKO8 cells transfected with empty vector control (EV), wildtype, or mutant DNMT3L. Data represent average values of the whole array from two independent experiments. Bottom, western blot analysis of DNMT3L protein expression in each cell line. For gel source data, see **Supplementary Fig. 1**. **f,** Methylation heatmap of 217,908 probes showing a difference in β-value greater than 0.3 in wildtype or mutant DNMT3L compared to those in empty vector control. Each row represents one CpG probe. **g**, Box plots of the 217,908 probes, showing the distribution of DNA methylation levels for each cell line. Median and interquartile range are represented by the bar and box. Dashed lines, background median β-value in DKO8 cells (top) and in wild-type DNMT3L-transfected DKO8 cells (bottom).

Moreover, by overlaying the recently resolved DNMT3A1 ubiquitin-dependent recruitment (UDR) maps^28–30^ onto our DNMT3A2-3L-nucleosome map, we found that the unoccupied acidic patch due to the 180° rotated conformation of the DNMT3L C terminal “Switching Helix” could potentially accommodate the DNMT3A1 UDR motif (**Fig. 4d**). This suggests that the full-length DNTM3A1, in the presence of DNMT3L, may adopt a similar conformation when binding to nucleosomes and function in a similar dynamic DNA engagement model.

### DNMT3L C-terminal “Switching Helix” and ADD domain are important for DNA methylation in cells

To assess the role of DNMT3L C-terminal “Switching Helix” and ADD domain in restoring DNA methylation in cells, we analyzed DNA methylation in DKO8 cells expressing either wild-type or mutant DNMT3L 56 days post-infection using the Infinium MethylationEPIC BeadChip array. Consistent with previous reports^4^, wild-type DNMT3L effectively restored DNA methylation to a relative high level (from β-value of 0.43 in the empty vector control to 0.71) (**Fig. 4e**). The E103R and Δα-helix complexes, which showed attenuated nucleosome binding, exhibited slightly reduced DNA methylation compared to wild-type DNMT3L complex with β-values of 0.69 and 0.67, respectively. In addition, the DNMT3L ADD deletion complex, which retained only ∼10% of nucleosome binding capacity, was much less efficient (with a β-value of 0.64) despite its elevated protein levels. Notably, the Δβ-sheet complex, while maintaining nucleosome binding capacity, showed the lowest levels of DNA methylation in cells (with a β-value of 0.49), likely due to insufficient protein expression (**Fig. 4e-f**). These results emphasize the importance of nucleosome recruitment and DNMT3L protein stability in DNA methylation in cells.

### DNMT3A2-3L senses histone tail and co-regulates DNA methylation

Although the histone tails beyond the nucleosome core were unresolved in the cryo-EM map, prior studies have showed that DNMT3A complexes interact with histone tails to target methylation to specific genome regions^9,13,18^. To explore this, we next used the AlphaScreen assay to assess the binding profiles for the whole panel of DNMT3A2-related complexes. Both DNMT3A2-3B3 and DNMT3A2-3L complexes showed strong nucleosome binding (**Fig. 5a-b**). Surprisingly, these two complexes displayed distinct behaviors depending on different histone modifications. While DNMT3A2-3B3 bound uniformly to nucleosomes regardless of histone modifications, DNMT3A2-3L nucleosome binding was modulated by histone modifications. Specifically, nucleosomes with H3K4me3, typically associated with active promoters and low DNA methylation^31^, inhibited the binding, whereas nucleosomes carrying H3K36me2/3, which are associated with gene body DNA methylation^32,33^, exhibited enhanced binding. Further analysis revealed that DNMT3A2 alone bound moderately to H3K36-methylated nucleosomes, while DNMT3L alone showed only very weak binding, suggesting a regulatory role for DNMT3L (**Fig. 5b**). Additionally, the binding profile of DNMT3B3 mirrored that of DNMT3A2-3B3 (**Fig. 5b** and **Extended Data Fig. 6**), indicating that DNMT3B3 might facilitate DNMT3A2-3B3 binding to a broader range of nucleosomes through acidic patch interactions to ensure continuous *de novo* DNA methylation in somatic cells. In contrast, DNMT3L, predominantly expressed in embryonic cells where DNA methylation patterns are established, likely modulates DNMT3A2-3L complexes to sense histone modifications and co-regulate DNA methylation. Interestingly, the methylation status of linker DNA selectively affected the binding of DNMT3A2-3L to nucleosomes, further suggesting special regulatory roles for the DNMT3L accessory protein (**Fig. 5c**).

**Fig. 5.**
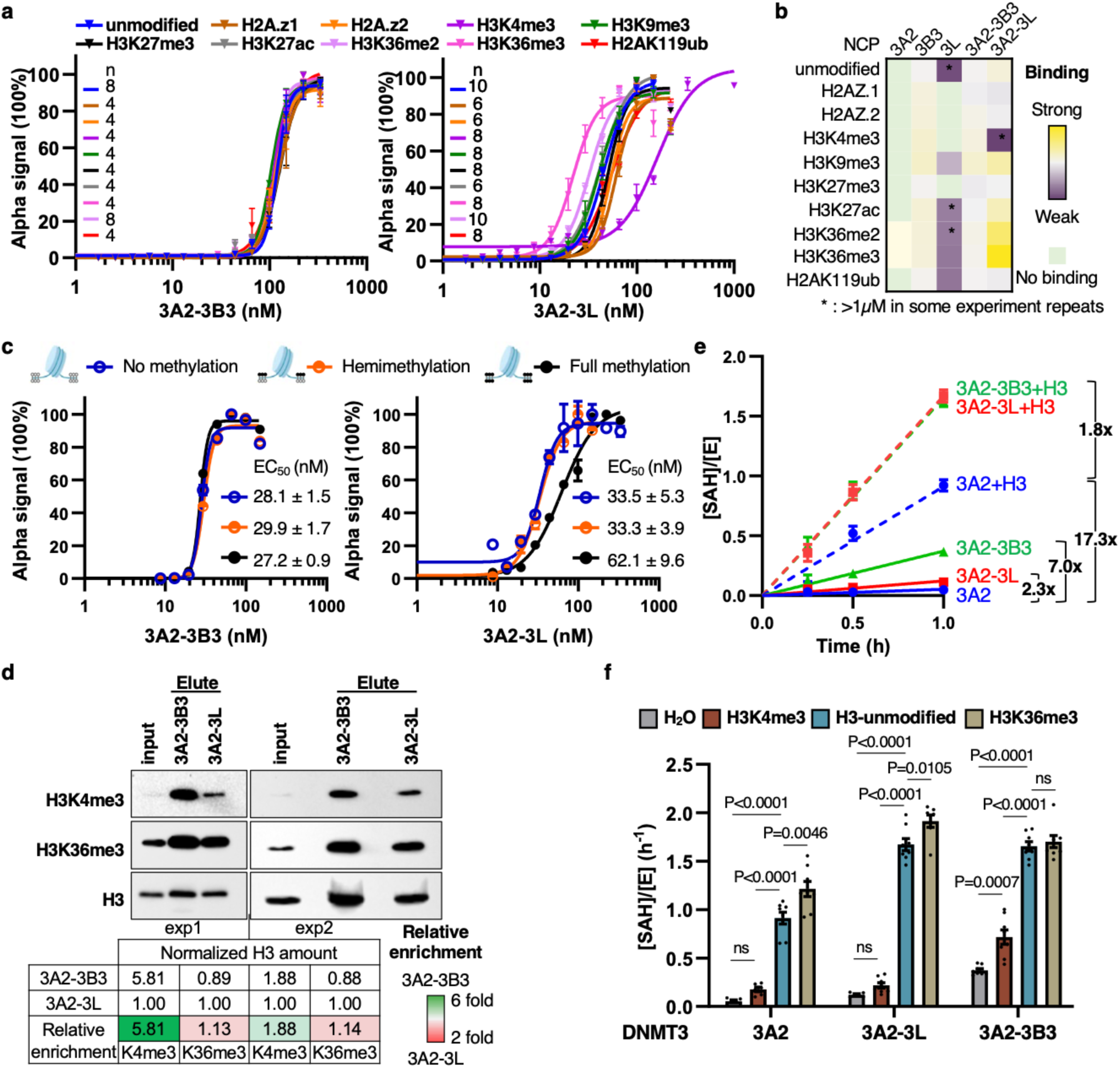
DNMT3A2-3L senses histone tail and co-regulates DNA methylation. **a,** AlphaScreen assay for the interaction of His-tagged DNMT3A2-3B3 (left) and His-tagged DNMT3A2-3L (right) complexes titrated against modified nucleosomes. Data were normalized within each group to the highest value (set as 100%). **b,** The full panel of DNMT3A2 nucleosome binding in AlphaScreen assay. The EC_50_s were shown in **Extended Data Fig. 6b**. **c,** Binding affinity curves and EC_50_s of DNMT3A2-3B3 and DNMT3A2-3L for linker DNA methylated nucleosomes. **d,** Western blot analysis of biotinylated protein complex pulldowns of native nucleosomes. For gel source data, see **Supplementary Fig. 1**. **e,** Free DNA methylation activities of DNMT3A2 with/without accessory proteins and histone H3 N terminal (1-44) peptides. **f,** Grouped DNA methylation activities of DNMT3A2, DNMT3A2-3L, DNMT3A2-3B3 with/without histone H3 N terminal (1-44) peptides unmodified or carrying K4 or K36 tri-methylation. Data were presented as mean ± s.e.m., n = 4-10 (**a**), 2 (**c**), and 8 (**e, f**). ns, not significant. P value, one-way ANOVA with Tukey’s multiple comparisons test.

To assess nucleosome binding in a more physiological context, we used biotinylated DNMT3A2-3B3 or DNMT3A2-3L complexes to pull down nucleosomes isolated from living cells. Although DNMT3A2-3B3 showed a higher overall pull-down efficiency (more H3), the trend remained consistent: DNMT3A2-3B3 exhibited higher efficiency (2-6 fold) in pulling down nucleosomes with H3K4me3 compared to DNMT3A2-3L, whereas DNMT3A2-3L showed slightly higher enrichment for nucleosomes with H3K36me3 (**Fig. 5d**). Notably, these effects were not as significant as those observed in the *in vitro* AlphaScreen assays, indicating a more complex regulation in living cells.

We next employed the MTase-Glo^TM^ methyltransferase assay to evaluate the DNA methylation activity of DNMT3A2-ralated complexes. As expected, DNMT3A2 alone showed minimal activity, while the accessory protein DNMT3L enhanced DNMT3A2 catalytic activity by approximately 2-fold, and DNMT3B3 increased it for an additional 3-fold, consistent with prior findings that heteromeric DNMT3A complexes are more stable and active^8,11,22–24^. Remarkably, Histone H3 peptide (1-44) increased DNMT3A2 catalytic activity by approximately 17-fold, far exceeding the stimulation observed with any accessory protein alone (2.3-fold and 7.0-fold). When combined with H3 peptides, accessory proteins further increased activity by another 2-fold (**Fig. 5e** and **Extended Data Fig. 7a-b**). These results aligned with in-cell data showing that DNMT3A2 overexpression restored genome-wide DNA methylation to a relatively high level (data not shown). In agreement with previous findings^5^, our data indicated that the gain in DNMT3A2 catalytic efficiency by accessory proteins and histone H3 peptides is primarily driven by an improved catalytic rate (K_cat_): in the absence of H3 peptide, DNMT3L enhanced K_cat_ of DNMT3A2 by 10-fold (0.32 h^-1^ vs 0.03 h^-1^), and DNMT3B3 by 30-fold (0.96 h^-1^vs 0.03 h^-1^); H3 peptide itself increased the catalytic rate more than 50-fold (1.77 h^-1^ vs 0.03 h^-1^), with further stimulation by accessory proteins (**Fig. 5e** and **Extended Data Fig. 7c**). In contrast, DNA substrate binding (K_m_) remained relatively stable, ranging from 0.42 to 0.83 µM in the absence of H3 peptide and 0.54 to 1.03 µM in the presence of H3 peptide (**Extended Data Fig. 7c**), suggesting that the binding and catalysis are not strictly correlated. Importantly, DNMT3A2-3L mutants that affect nucleosome binding did not reduce the methylation of naked DNA *in vitro* (**Extended Data Fig. 7d**), supporting our hypothesis that DNMT3 binding to nucleosomes and methylating DNA are two relatively separate biological processes, potentially regulated by different factors. Notably, our in-cell DNA methylation data and free DNA catalytic analysis suggest that the observed defects in CpG methylation in cells were due to defects in nucleosome targeting, not DNA binding or catalysis.

In cells, DNA methylation is regulated by chromatin environment, including multiple post-translational modifications of histones, such as methylation at H3K4 and H3K36. To investigate this, we tested the stimulation capacity of histone H3 peptides harboring K4me3 and/or K36me3, as these modifications significantly altered the nucleosome binding behavior of the DNMT3A2-3L complex. As expected, H3K4me3-modified peptides diminished the stimulation capacity, even in the presence of accessory proteins. Notably, DNMT3A2-3B3 complex still retained relatively high activity in the presence of H3K4me3, probably due to the high basal activity of this complex (**Fig. 5f**). Interestingly, H3K36me3 slightly enhanced activity of 3A2 and 3A2-3L, and could partially attenuate the inhibitory effect of H3K4me3 (K_cat_ from 0.83 h^-1^ to 1.63 h^-1^) – but only in the DNMT3A2-3L complex, not in DNMT3A2-3B3 (**Fig. 5f** and **Extended Data Fig. 7a-b, e**), indicating the histone modification regulation is complicated and environment dependent. By further investigating H3 peptide, we identified that the N-terminal region (residues 1-21), which includes K4, retained stimulatory potential (**Extended Data Fig. 7e**), which is consistent with the previous experiments focused on DNMT3A2^34^.

Based on these findings, we propose that while accessory proteins are important for DNMT3A2 activity, histone H3 may play a more direct stimulatory role *in vivo*. Meanwhile, accessory proteins also have regulatory functions on DNA methylation beyond activity stimulation. Further studies will be needed to map functional regions of histone H3 and further elucidate the regulatory roles of histone modifications in DNA methylation in living cells.

### DNMT3A2-3L dynamically searches for DNA and engages with nucleosomes

The distal DNMT3A2 CD domain exhibits weak DNA binding (**Extended Data Fig. 4g**), suggesting a flexible interaction between DNMT3A2-3L and DNA. To explore the dynamics of the DNMT3A2-3L-NCP complex, we performed 3D classification without alignment in RELION5, which revealed a swing motion in the distal region of the DNMT3A2-3L complex (**Extended Data Fig. 1e**). This finding was confirmed by the recently developed CryoROLE (CryoEM Relative Orientation LandscapE)^35^. Four representative centered points were selected and superimposition of the reconstructed maps from each point within 5 Å revealed “up” and “down” movements of the distal DNMT3A2-3L (**Extended Data Fig. 8a-c**). For a more detailed analysis of the relative orientations between DNMT3A2-3L and nucleosomes, 3D reconstructions were calculated from locations at every 10° along each direction. The landscape analysis identified swiveling motion in the α direction and swinging motion in the β and γ directions (**Extended Data Fig. 8d** and **Supplementary Video 1**). These movements were also captured by 3D multi-body refinement in RELION5 within the same dataset (**Extended Data Fig. 8e** and **Supplementary Video 2**). These findings underscore the dynamics of the DNMT3A2-3L-NCP complex, especially the distal DNMT3A2-3L region and the linker DNA. With an additional cryo-EM dataset, we identified three different conformational states of the nucleosome-bound DNMT3A2-3L complex, which recaptured the swing motion: a DNA-engaged state, an intermediate state, and an “up” state (**Fig. 6a** and **Extended Data Fig. 2**).

**Fig. 6.**
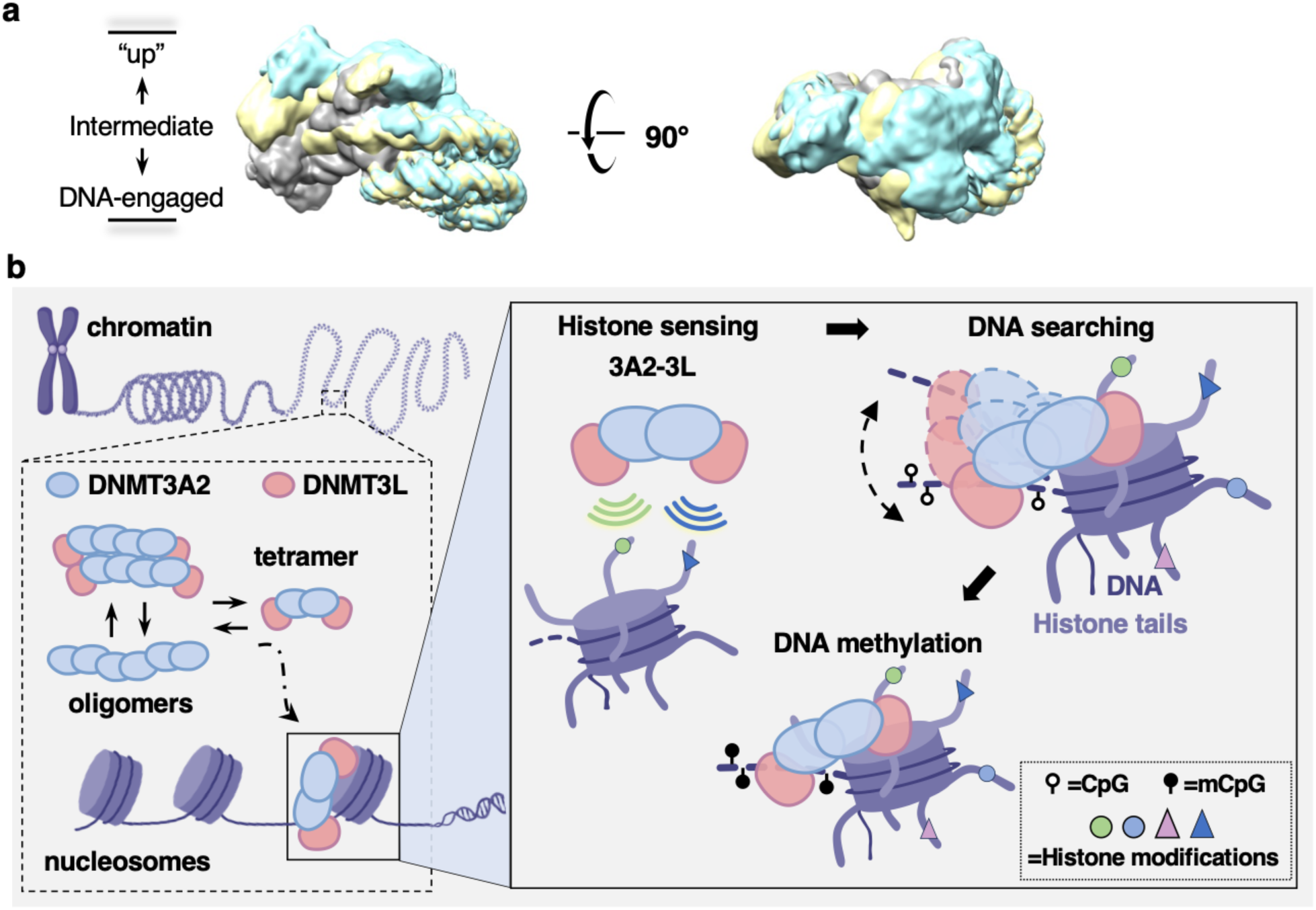
Model for DNMT3A2-3L-mediated *de novo* DNA methylation on chromatin. **a,** Cryo-EM data analysis revealed three different conformational states of the DNMT3A2-3L-NCP complex: a DNA-engaged state (grey), an intermediate state (yellow), and an “up” state (cyan). **b,** A proposed model for DNMT3A2-3L-mediated *de novo* DNA methylation on chromatin by complex reorganization, histone sensing, dynamic DNA search and nucleosome engagement.

Based on these observations and biochemical data, we propose a model for DNMT3A2-3L-mediated *de novo* DNA methylation on chromatin in which DNMT3A2-3L re-organizes its oligomeric states, senses histone tails, and dynamically interacts with nucleosomes to achieve optimal DNA engagement for methylation (**Fig. 6b**). Even in the presence of the accessory protein DNMT3L, DNMT3A2 prefers to form higher-order oligomers such as the dodecamer. Upon binding to nucleosomes, it transitions into a more stable and active heterotetrameric conformation, which can sense histone modifications to cooperate with other epigenetic regulations and dynamically search for DNA engagement through interactions between DNMT3L and nucleosome cores. These flexible interactions, potentially involving charge and polar interactions, could be regulated by histone modifications. Such flexibility allows the distal part of DNMT3A2-3L to move “up” and “down”, facilitating engagement with linker DNA for methylation.

## Discussion

The oligomeric assembly of DNMT3 complexes plays a crucial role in DNA methylation^36^. While the heterotetrameric complex formed by the isolated C-terminal domains of DNMT3A, DNMT3B, and DNMT3L has been well-characterized^8–10,25^, the mechanism of the DNMT3A-3L complex binding to nucleosomes remained unclear. Our study addresses this knowledge gap by structural and biochemical characterization of the DNMT3A2-3L complex, both in its nucleosome-bound state and its nucleosome-free state. Our findings provide key insights into its dynamic assembly on nucleosomes and molecular mechanisms of *de novo* DNA methylation on chromatin.

First, our cryo-EM structure of the DNMT3A2-3L-nucleosome complex revealed a domain organization – asymmetrically binding to nucleosomes – that closely resembles the previously reported DNMT3A2-3B3-nucleosome structure^10^. Specifically, the proximal DNMT3L subunit interacts with the nucleosome core, stabilizing the complex and facilitating recognition of the nucleosome core surface, while the distal DNMT3A2 subunit binds to linker DNA, positioning the catalytic domain for precise DNA methylation. This asymmetric arrangement provides a crucial structural basis for the regulation of *de novo* DNMT3 complexes in early embryonic development. Interestingly, only one DNMT3A2 subunit actively engages with DNA, potentially explaining the clustering of CpG sites at variable distances from 1 to 25 bps in the human genome^37^, rather than the 12-bp intervals observed in the heterotetramer of CD and CLD domains of DNMT3A and DNMT3L^6,8^. DNMT3A tends to form large oligomers through self-association along its R-D and F-F binding surfaces^24,26,38^, whereas DNMT3L can only form the F-F interfaces, preventing further oligomerization and resulting in a DNMT3L-3A-3A-3L tetramer of the C-terminal domains^26,39^. Consistent with the literature, our data also showed that DNMT3L occupied F-F interfaces, resulting in a DNMT3L-3A2-3A2-3L heterotetramer in its nucleosome-bound state. Additionally, while recent DNMT3B homo-oligomerization has been described^27^, our study reveals, for the first time, a novel higher-oligomeric arrangement of DNMT3A2-3L complex in its nucleosome-free state. This suggests its dynamic reorganization and allostery regulation of the DNMT3A2-3L complex upon binding to nucleosomes by subunit architecture reorganization. The biological function of this hetero-oligomeric state remains an open question. Further exploration is needed to understand its potential roles in DNA methylation and its implications for genomic regulation during development.

Second, this study provides new insight into the distinct role of DNMT3L as an accessory protein. In somatic cells, where DNA methylation patterns are already established, *de novo* DNMT3 enzymes cooperate with DNMT3B3 for continuous *de novo* DNA methylation due to DNA methylation loss during replication and DNA repair^40,41^. In contrast, DNMT3L is largely expressed in embryonic cells, where DNA methylation patterns are about to be established. This cell-specific expression suggests distinct roles for each accessory protein. Our cryo-EM structure and nucleosome-binding assays highlight a unique DNMT3L nucleosome binding mechanism. Specifically, DNMT3L anchors to the nucleosome core through its ADD domain and C-terminal “Switching Helix”. This binding mode is distinct from DNMT3B3 using R-fingers as demonstrated in our previous study^10^. This interaction, potentially mediated by moderate-range charge and polar interactions, contributes to the flexibility of active subunits to facilitate the engagement with linker DNA, as discussed below. The binding differences were also confirmed by the native nucleosome pulldown experiment, supporting the notion that the DNMT3A2-3L complex senses the histone modifications in the cellular environment. Furthermore, aligning our map with recently identified DNMT3A1 N-terminal UDR motif^28–30^, we observed that the 180-degree rotated conformation of the C-terminal “Switching Helix” in DNMT3L can perfectly accommodate the UDR density and provide sufficient space for the PWWP domains, even though they were not observed in our current cryo-EM map. This consistency further indicates that DNMT3L, as an accessory protein, utilizes a similar mechanism in the nucleosome binding of the DNMT3A1-3L complex.

Third, our nucleosome-binding assays and DNA methylation assays indicated that DNMT3A complex binding to nucleosomes may not be directly coupled with DNA methylation activity. These two biological events, while interconnected, could be independently regulated by different factors such as histone modifications. Additionally, although accessory proteins can stimulate DNMT3A catalytic activity^4,22,23,42^, histone tails appear more potent, and the combination of these two shows the highest capacity. This highlight the role of accessory protein not only in stimulating activity but also in regulating complex architectures coordinating DNA methylation with other biological processes.

Finally, our dynamic and flexibility analyses suggested a “search-and-engage” model for the active subunit (DNMT3A2) to interact with a linker DNA by conformational changes. Nucleosome binding may release the higher-oligomeric state into a more stable and active heterotetrameric state of DNMT3A2-3L, in which the complex can sense the histone modifications to cooperate with other epigenetic regulations and anchor to the nucleosome core through the DNMT3L ADD domain and C-terminal “Switching Helix”. This anchoring is not stiff, allowing the active DNMT3A2 subunit to seek an optimal DNA engagement position for linker DNA methylation. While this study primarily focused on the DNMT3A2-3L complex, we hypothesize that DNMT3L may function similarly in the DNMT3B or DNMT3A1 complex. *In vivo*, most CpG sites are methylated without strict spacing patterns, implying the dynamic regulation of DNMT3 in complex with DNMT3L.

In summary, our structural and functional analysis revealed a novel mechanism by which DNMT3A-3L mediates de novo DNA methylation on chromatin through complex reorganization, histone tail sensing, dynamic DNA search, and nucleosome engagement. This provides invaluable mechanistic insights into how DNA methylation is established and regulated on chromatin, laying the foundation for exploring new therapeutic strategies targeting aberrant DNA methylation.

## Acknowledgments

We thank G. Zhao and X. Meng from Van Andel Institute (VAI) for the support with data collection at the David Van Andel Advanced Cryo-Electron Microscopy Suite; and VAI Genomics Core for sequencing and Illumina Infinium methylation EPIC array support. We are grateful to W. Choi, C. Li, and Y. Cheng from the University of California San Francisco (UCSF) for providing the CryoROLE algorithms and assisting in the use of the algorithms; J.C. Eissenberg and E. Di Cera from Saint Louis University (SLU), and B.M. Dickson (VAI) for advice.

This work was supported by the Doisy Fund of the Edward A. Doisy Department of Biochemistry and Molecular Biology at Saint Louis University School of Medicine (T.H.X.), NIH R35CA209859 (P.A.J.), NIH R35GM147261 (E.J.W.), and NIH R50CA243878 (M.L.).

## Author contributions

T.H.X. and P.A.J. conceived the project. Y.Y. and T.H.X. prepared protein samples, performed EM experiments and data analysis. Y.Y. and T.H.X. performed and interpreted all experiments. Y.Y., T.H.X., and X.E.Z. built the atomic models and refined the atomic models. M.L. performed DKO8 cell transfection and methylation EPIC array and data analysis. S.L.T. assisted in experiments. G.L. assisted in DNA cloning. E.J.W. supervised X.E.Z.; P.A.J. supervised S.L.T. and M.L.; T.H.X. supervised Y.Y. and G.L.. Y.Y. and T.H.X. wrote the paper with support from all authors, and the manuscript has been reviewed and approved by all authors.

## Data availability

The 3D cryo-EM maps have been deposited in the Electron Microscopy Database under accession numbers EMD-48322 (DNMT3A2-3L-NCP consensus map, 3.10 Å), EMD-48492 (DNMT3A2-3L, 3.60 Å), EMD-48493 (NCP, 3.08 Å), EMD-48495 (DNMT3A2-3L-NCP intermediate state map, 3.29 Å), EMD-48494 (DNMT3A2-3L-NCP “up” state map, 3.16 Å), EMD-48498 (combined DNMT3A2-3L-NCP map, 3.60 ∼ 3.08 Å), EMD-48496 (DNMT3A2-3L dodecamer, 3.66 Å), and EMD-48497 (DNMT3A2 hexamer, 3.62 Å). The structure coordinates have been deposited in the Protein Data Bank under accession numbers PDB ID 9MPP (DNMT3A2-3L-NCP), PDB ID 9MP0 (DNMT3A2-3L dodecamer), and PDB ID 9MPO (DNMT3A2 hexamer). The DNA methylation data have been deposited in the GEO database with accession code GSE291793. All other data needed to evaluate the conclusions in the paper are presented in the main text or supplementary materials. No restrictions are placed on data availability. Source data are provided with this paper.

## Methods

### Cell culture

Sf9 insect cells were obtained from Expression Systems and grown in ESF-921 medium (Expression Systems) at 110 rpm, 27 °C.

HEK293T cells were obtained from the American Type Culture Collection and cultured in Dulbecco modified Eagle medium (HyClone) supplemented with 10% fetal bovine serum (FBS, Corning). The DKO8 cells, obtained from Dr. Stephen Baylin’s laboratory at Johns Hopkins University, were cultured in McCoy’s 5A medium (Gibco) containing 10% FBS and 1% penicillin/streptomycin. All cells were maintained at 37°C in a humidified incubator with 5% CO_2_ and routinely verified to be free of *Mycoplasma* contamination.

### DNA cloning and protein expression

*Xenopus laevis* histone genes were from plasmid pET29a-YS14 (kind gift of Dr. Jung-Hyun Min, Washington State University). Full-length human DNMT3A and DNMT3L genes were purchased from DNASU (HsCD00620811, HsCD00077165). All site-directed mutagenesis was carried out using the PCR method. All constructs were confirmed by DNA sequencing. DNA and protein preparation is followed by the previous protocol^10^.

Briefly, *E. coli* BL21(DE3) pLysS cells transformed with two His6-Sumo tags *Xenopus laevis* core histones polycistronic expression plasmids were grown in LB medium to an OD_600_ of ∼1 at 37 °C and induced with 500 μM isopropyl-β-Dthio-galactopyranoside (IPTG) at 37 °C for 4-5 h. Cells were harvested, resuspended, and lysed in 20 mM Tris-HCl pH 8.0, 2.0 M NaCl, 25 mM imidazole, 10% glycerol, 1 mM phenylmethanesulfonyl fluoride (PMSF), and 0.5 mM tris(2-carboxyethyl)phosphine (TCEP), using a high-pressure homogenizer (APV). Lysates were cleared by centrifugation for 45 min at 38,700 x g at 4 °C, passed over a 5-mL HisTrap FF column (Cytiva), and eluted with 20 mM Tris-HCl pH 8.0, 2.0 M NaCl, 250 mM imidazole, 10% glycerol, and 0.5 mM TCEP. His6-Sumo tags were cleaved by ULP1 Sumo protease and subsequently purified by size-exclusion chromatography (SEC) through a HiLoad 26/60 Superdex 200 column (Cytiva) in 10 mM Tris-HCl pH 8.0, 2.0 M NaCl, 1 mM EDTA, and 0.5 mM TCEP. The histone octamer peak fractions were pulled, verified by 18% SDS-PAGE, concentrated, and fresh-frozen in aliquots in the presence of 50% glycerol for storage.

The His8-GFP tagged and untagged human DNMT3A2 WT or mutants and human DNMT3L WT or mutants were cloned into pFastBac (ThermoFisher) vectors and expressed in Sf9 insect cells using the Bac-to-Bac system (ThermoFisher). The insect cells were harvested, resuspended, and lysed in 20 mM Tris-HCl pH 8.0, 300 mM NaCl, 25 mM imidazole, 50 µM ZnSO_4_, 10% glycerol, 0.03% 2-mercaptoethanol, 1 mM PMSF, and 1x EDTA-free protease inhibitor cocktail (Roche) using Dounce homogenization. Lysates were centrifuged at 150,000 x g at 4 °C for 1 h and the supernatant was passed through a 5-mL HisTrap FF column (Cytiva), and eluted with 20 mM Tris-HCl pH 8.0, 300 mM NaCl, 250 mM imidazole, 50 µM ZnSO_4_, 10% glycerol, and 0.03% β-mercaptoethanol. The elute was subsequently purified by SEC through a HiLoad 16/600 Superdex 200 pg column (Cytiva) in 20 mM HEPES pH 8.0, 300 mM NaCl, 50 µM ZnSO4, 10% glycerol, and 3 mM DTT or digested with purified TEV protease at 4 °C overnight to remove the His8-GFP tag. The TEV-digested DNMT complex was further purified by SEC. The His8-GFP tagged and untagged DNMT complex peak fractions were pulled, verified by 12.5% SDS-PAGE, concentrated, and fresh-frozen in aliquots for storage.

### Nucleosome DNA template preparation

The nucleosome DNA template designs are based on the 147-bp Widom 601 nucleosomal positioning sequence^21^ with the 10 bp linkers that we identified previously^10^ at the entry and exit sites, generating Widom601-167 for cryo-EM study, and 301-bp DNA template (Widom601-301)^10,43^ for the DNA methylation study. The sequences for Widom601-167 are: 5’-ATCGGCCGCC**CTGGAGAATCCCGGTGCCGAGGCCGCTCAATTGGTCGTAGACAGCTCTA GCACCGCTTAAACGCACGTACGCGCTGTCCCCCGCGTTTTAACCGCCAAGGGGATTACTCCCTAGTCTCCAGGCACGTGTCAGATATATACATCCTGT**GGCGGCCGAT-3’, for Widom601-301 are: 5’-GATAGACAGCTGCTGAACCAATGGGACCAAGCTTCACACCGAGTTCATCGCTTATGTGATC GACCATCGGCCGCCTA**CTGGAGAATCCCGGTGCCGAGGCCGCTCAATTGGTCGTAGACA GCTCTAGCACCGCTTAAACGCACGTACGCGCTGTCCCCCGCGTTTTAACCGCCAAGGGG ATTACTCCCTAGTCTCCAGGCACGTGTCAGATATATACATCCTGT**CAGGCGGCCGATTGTAT TGAACAGCGACCTTGCCGGTGCCAGTCGGATAGTGTTCCGAAAGCTTCTGCCCAACTGGC.

The 601 sequence is shown in bold. The 12x Widom601-167 was expressed in *Escherichia coli* using pUC19 plasmid and isolated as described previously^10^. The Widom601-301 were amplified using polymerase chain reaction (PCR) using an appropriate primer pair (301F: GATAGACAGCTGCTGAACCAATGGG and 301R: GCCAGTTGGGCAGAAGCTTTC). Phusion polymerase was used to amplify 10 mL reaction volumes in a 96-well plate. PCR products were purified on a 6-mL anion exchange chromatography column (Cytiva), precipitated with ethanol, and dissolved in ddH_2_O.

### Nucleosome preparation

Optimized ratios of octamer to DNA were mixed and nucleosomes were reconstituted by the double-dialysis bag method^10^ or stepwise salt dialysis against low salt buffers to a final buffer containing 20 mM HEPES, pH 8.0, 50 mM KCl, 1 mM EDTA, and 1 mM DTT. Reconstituted nucleosomes were analyzed on 6% PAGE.

For *in vitro* nucleosome binding AlphaScreen assay, the biotinylated nucleosomes harboring different histone tail modifications (bio-NCP147) and linker DNA methylation statuses (bio-NCP199) were purchased from Epicypher.

### DNMT-nucleosome complex formation

For cryo-EM studies, 1.5 µM DNMT3A2-3L complex was incubated with 1 µM NCP167 for 30 min on ice in 20 mM HEPES pH 8.0, 50 mM NaCl, 1 mM MgCl_2_, 1 mM DTT, and 100 µM SAH, and was then loaded onto a 12-mL linear 10%-25% (v/v) glycerol gradient in the same buffer supplemented with 0%-0.15% EM-grade glutaraldehyde^44,45^. After centrifugation at 4 °C for 18 h at 38,000 rpm in an SW41 rotor (Beckman Coulter), the sample was fractionated from the bottom to the top. Fractions were analyzed by native PAGE, and protein complex-containing fractions were pooled and buffer exchanged with 20 mM HEPES pH 8.0, 50 mM NaCl, 1 mM MgCl_2_, 1 mM DTT, and concentrated to approximately 0.2 mg/mL.

### Cryo grid preparation

A droplet (3.0 µL) of cross-linked DNMT-NCP complex at a concentration of about 0.2 mg/mL was applied to a glow-discharged holey carbon grid (Quantifoil R1.2/1.3, Au 300 mesh), and subsequently blotted for 3 s and vitrified by plunging into liquid ethane with a Vitrobot Mark IV (Thermo Fisher Scientific) operated at 4 °C and 100% humidity. Each grid was screened using 200 keV Talos Arctica before data collection.

### Cryo-EM data acquisition, processing and 3D reconstructions

Cryo-EM data were collected on a Titan Krios transmission electron microscope (TEM) operated at 300 keV and equipped with a K3 summit direct detector (Gatan). Automated data acquisition was carried out using Serial EM^46^ in super-resolution mode at a magnified pixel size of 0.414 Å, with defocus values ranging from −1.0 to −2.0 µm. The total exposure time was set to 2 s with 65 frames, resulting in an accumulated dose of about 60 e^-^ per Å^2^. A total of 17,684 image stacks (data1) and 14,303 image stacks (data2) were collected for the DNMT3A2-3L-NCP complex, and 16,686 image stacks were collected for the DNMT3A2-3L oligomer complex.

The image stacks were processed using cryoSPARC Live (v4.6.0), where patch-based motion correction was performed with 2× binning (0.828 Å/pixel) and patch-based contrast transfer function (CTF) was estimated, followed by manual inspection to remove images that contained contaminated crystalline ice, other forms of visible contamination, or that were taken from broken holes. Micrographs with an estimated maximum resolution beyond 5 Å were discarded, and high-quality, motion-corrected sums were used for image processing. Global and local resolution estimates were calculated in cryoSPARC (v4.6.0) using the gold-standard Fourier shell correlation criterion. Final cryo-EM maps were sharpened by DeepEMhancer^47^.

For the DNMT3A2-3L-NCP dataset1, high-quality micrographs were selected for auto picking and particle extraction. Approximately 5 million particles were initially extracted at 4x-downscaling and subjected to reference-free 2D classification. About 1 million good particles were selected for Ab-Initio Reconstruction into six classes, of which five good ones were selected for further analysis. These selected particles were then re-extracted with a box size of 400 pixels without downscaling and Ab-Initio Reconstruction of these particles generated three good classes (∼65% of the total), which were subsequently refined with Homogeneous and Non-uniform Refinement, achieving a final global nominal resolution of approximately 3.14 Å. The homogeneous DNMT3A2-3L-NCP complex particles (627,284) were then imported into RELION (v5) for 3D classification without alignment and for DNMT and NCP particle subtraction. The subtracted particles were further processed in cryoSPARC (v4.6.0) to achieve the overall resolution at 3.08 Å for the NCP and 3.70 Å for the DNMT3A2-3L complex. These particles were also analyzed by Orientation Relation Landscape algorithms^35^ to virtualize the relative orientation between the DNMT and NCP (details provided in the Orientation Relation Landscape section). Local resolution estimates were determined by cryoSPARC (v4.6.0) at gold-standard 0.143 FSC. (**Extended Data Fig. 1**).

About 921,860 high-quality particles were selected by reference-free 2D and Ab-Initio Reconstruction from DNMT3A2-3L-NCP dataset2 and re-extracted with a box size of 400 pixels and combined with 627,284 high-quality particles of the same box size from DNMT3A2-3L-NCP dataset1 for further data processing. The Ab-Initio Reconstruction of the combined particles generated three distinct conformations: a DNA-engaged state (grey), an intermediate state (yellow), and an “up” state (cyan). Subsequent Heterogeneous, Homogeneous, and Non-uniform Refinement achieved an overall resolution of 3.10 Å for the engaged state, 3.29 Å for the intermediate state, and 3.16 Å for the “up” state. Particles from each conformation were further subtracted for DNMT and NCP for focused refinement. The overall NCP resolutions were achieved at 3.15 Å for the engaged state, 3.28 Å for the intermediate state, and 3.15 Å for the “up” state. The overall DNMT resolutions were achieved at 3.60 Å for the engaged state, 5.68 Å for the intermediate state, and 4.22 Å for the “up” state. Local resolution estimates were determined by cryoSPARC (v4.6.0) at 0.143 FSC (**Extended Data Fig. 2**).

For the DNMT3A2-3L oligomer complex, approximately 2 million particles were initially extracted at 4x-downscaling from selected high-quality micrographs and subjected to reference-free 2D classification and Ab-Initio Reconstruction into three classes, resulting in two good classes (∼75%): dodecamer and hexamer, for further analysis. The final global nominal resolutions were achieved at 3.66 Å for the dodecamer state and 3.62 Å for the hexamer state. Local resolution estimates were determined by cryoSPARC (v4.6.0) at 0.143 FSC (**Extended Data Fig. 3**).

### Model building and refinement

We utilized the composite map generated from focused refinements based on the consensus refinement model to provide better density for the model building of DNMT3A2-3L. Given the varying resolution limits across different DNMT regions, the whole DNMT3A2-3L-NCP model was generated by combining AlphaFold2^48,49^ predicted models and the previously published structures^6,10^, which were rigidly docked into the composite map. Specifically, the previously resolved DNMT3A2-3B3-NCP structure (PDB ID 6PA7) and the AlphaFold2 predicted DNMT3A2-3L complex were fitted into the cryo-EM density in ChimeraX^50^. All models were then iteratively manually adjusted in COOT^51^ and subjected to real-space refinement using PHENIX^52^. The model validation was performed with Molprobity in PHENIX^52^. Structural figures were prepared in ChimeraX^50^ and PyMOL (https://pymol.org/2/). The final refinement statistics were summarized in **Extended Data Table 1**. The extent of any model overfitting during refinement was measured by refining the final model against one of the half-maps and by comparing the resulting map versus model FSC curves with the two half-maps and the full model.

### Orientation Relation Landscape Analysis

Subtracted particles corresponding to DNMT and NCP from dataset1 were refined using manually curated DNMT region and NCP region maps to generate the particle-refined star files for Orientation Relation Landscape analysis^35^. Briefly, the OrientationRelation.py tool was employed to calculate the orientation relations between DNMT and NCP. Interactive 2D projection views of these orientations were generated using the Display_2D_Projections.py tool. For improved resolution, the parameter t was set to 1.2. Four representative centered points (**Extended Data Fig. 8c**) were selected. For each direction (α, β, γ), the selected particle coordinates were shown in **Extended Data Fig. 8d.** The particles within 5 Å were selected for 15 Å low-pass filter 3D reconstruction using the relion_reconstruct tool. The reconstructed maps from each region were combined based on the consensus-refined map.

### DNMT-nucleosome binding assay

The luminescence proximity AlphaScreen assay was used to assess the *in vitro* interactions between DNMT proteins and nucleosomes. For single-point experiments, 20 nM biotinylated nucleosomes and 100 nM His-tagged DNMT proteins were incubated with 5 µg/mL streptavidin-coated donor beads and 5 µg/mL nickel-chelated acceptor beads (Revvity) in 100 µL AlphaScreen buffer (50 mM MOPS, pH7.4, 50 mM NaF, 50 µM CHAPS, and 0.1 mg/ml BSA) for 1.5 h in the dark at room temperature. Photon counts were determined in 384 plates using an Envision-Alpha Reader (PerkinElmer). For AlphaScreen competition assays, 20 nM biotinylated nucleosomes and 100 nM His-tagged DNMT proteins were incubated in 100 µL AlphaScreen buffer containing 5 µg/mL of donor and acceptor beads each in the presence of 1 µM peptide competitors or 500 nM untagged DNMT3 complexes. For binding strength (EC_50_) assessment, 20 nM biotinylated nucleosomes were incubated with increased concentrations of His-tagged DNMT proteins in 100 µL AlphaScreen buffer containing 5 µg/mL of donor and acceptor beads each. The curves were plotted using photon counts (y-axis) against DNMT concentration (x-axis) and the EC_50_s were calculated by dose-response - stimulation with Hill slope using GraphPad Prism.

### DNA methylation kinetic assay

DNMT activity was measured by incorporating methyl groups from S-adenosylmethionine (SAM) into Widom601-301 DNA, using the MTase-Glo™ assay (Promega) following the manufacturer’s protocol. Briefly, 500 nM Wimdon601-301 DNA (12 µM CpG sites) was methylated in the presence of 200 nM purified DNMT complexes (0.4 µM catalytic sites) in 20 mM HEPES pH7.4, 50 mM NaCl, 5% glycerol, 1 mM MgCl_2_, 1 µM ZnSO_4_, 1 mM DTT, 0.1% BSA, and 10 µM SAM at 37 °C in 150 µL total volume with or without 2 µM histone H3 peptides. The reactions were incubated at 37 °C for variable time. At 0, 0.25, 0.5, 1, 2, and 4 h, 20 µL of reactions were taken and stopped by adding 5 µL of 0.5% TFA. A 17.5 µL of each stopped reaction was mixed with 3.5 µL of freshly prepared 6x MTase-Glo™ reagent and incubated at room temperature for 30 min, followed by another 30-min incubation with MTase-Glo™ detection solution at room temperature. Luminescence was measured in a 384-well plate with 12 µL in each well using an Envision-Alpha Reader (PerkinElmer). The standard curve for *S*-adenosyl homocysteine (SAH) was generated by serial dilution of SAH standard (150 µL each from 0 to 1.0 µM) and the luminescence measurement was performed exactly as the activity assay. For each standard: net RLU (luminescence) = RLU of the standard - RLU of the 0 µM SAH standard. For each sample: net RLU = RLU (with enzyme) - RLU (without enzyme). The standard curve was plotted using net RLU (y-axis) against SAH concentration (x-axis), and a linear regression graph was generated using GraphPad Prism. The linear equation was used to calculate the SAH concentration of each reaction sample. The kinetics measurements were performed for 1 h at 37°C with a total volume of 20 μL and varying concentrations of Widom601-301 DNA substrate from 0 to 500 nM. The reaction was stopped by adding 5 µL of 0.5% TFA and 17.5 µL stopped reactions were used for SAH measurement.

### Native nucleosome isolation

About 30 million HEK293T cells were resuspended in 500 µL ice-cold Buffer N (25 mM Tris-HCl pH 7.5, 15 mM NaCl, 60 mM KCl, 8.5% Sucrose, 5 mM MgCl_2_, 1 mM CaCl_2_, 200 µM PMSF, and 1x Protease inhibitor cocktail (Roche)) and lysed by adding equal volume of Buffer N with 0.6% NP-40 substitute on ice for 16 min. Nuclei were pelleted by centrifugation at 5,000 rpm 10 min at 4 °C and then washed twice with Buffer N. The washed nuclei were resuspended in 500 µL Buffer N, and an equal volume of sucrose cushion (25 mM Tris-HCl pH 7.5, 15 mM NaCl, 60 mM KCl, 30% Sucrose, 5 mM MgCl_2_, 1 mM CaCl_2_, 200 µM PMSF, and 1x Protease inhibitor cocktail [Roche]) was added. After incubating on ice for 3 min, nuclei were pelleted again by centrifugation at 5,000 rpm for 10 min at 4 °C and resuspended in 500 µL MNase digestion buffer (25 mM Tris-HCl pH 7.5, 5 mM MgCl2, 5 mM CaCl2, 8.5% Sucrose, 10% Glycerol, 200 uM PMSF). Chromatin DNA concentration was measured using a Nanodrop after taking 2 µL of nuclei and adding 18 µL of 2M NaCl. Micrococcal nuclease (MNase, New England Biolabs) was added at a concentration of 200 gel units per 10 µg nuclei, and digestion was carried out for 20 min at 37 °C. Reactions were terminated by adding 1/10 volume of 0.5M EDTA and nucleosomes were harvested by centrifugation at 5,000 rpm for 10 min at 4 °C to remove debris.

### Native nucleosome pull-down for histone modification binding analysis

Native nucleosomes (∼150 µg DNA) were incubated overnight with streptavidin MagBeads (Genescript) pre-bound to ∼70 pmol biotinylated DNMT3A2-3L or biotinylated DNMT3A2-3B3. After washing, bound proteins were eluted with 50 µL of SDS loading buffer. 10 µL of eluted protein from each sample was resolved on a 12.5% SDS-PAGE gel and transferred to a polyvinylidene difluoride (PVDF) membrane (Genescript). The membrane was blocked with 5% milk in TBST (20 mM Tris-HCl pH 7.4, 150 mM NaCl, and 0.1% Tween 20) and incubated overnight at 4 °C with antibodies against histone H3K36me3 (Cell Signaling Technology 9763S). After washing 3 times with TBST, membranes were incubated with an HRP-conjugated secondary antibody. Proteins were detected using ECL western blotting detection reagent (BioRad) and imaged with the iBright system (Thermo Fisher Scientific). To assess the histone H3K4me3, the membrane was stripped and re-probed with an antibody against histone H3K4me3 (Cell Signaling Technology 9751S). To assess total histone H3 levels, the membrane was stripped and re-probed with an antibody against histone H3 (Cell Signaling Technology 9715S). Each exposure was analyzed in iBright system (Thermo Fisher Scientific) using 3A2-3L pull-down within each group as a reference and normalized by total histone H3.

### Lentivirus generation and infection

Human DNMT3L and its mutants, tagged with an N-terminal Myc-tag were cloned into pLJM1 lentiviral vector according to the manufacturer’s protocol. The production of lentivirus and cell infection was performed as previously described^4,10^. Briefly, the lentiviral plasmid, along with the packaging plasmids pMD2.G and psPAX2 (Addgene plasmid #12259 and #12260), were co-transfected into HEK-293T cells using Lipofectamine LTX (Thermo Fisher Scientific) and OPTI-MEM media (Invitrogen). Lentiviruses were harvested twice on day 3 and day 5. Viruses were filtered through 0.45 μm filters and stored at −80 °C. To infect DKO8 cells, lentivirus was added to the culture media with 8 μg/mL polybrene (Sigma H9268). Forty-eight hours post-infection, cells were selected with 2 μg/mL puromycin until harvesting.

### Detection of DNMT3L protein levels

DKO8 cells carrying Myc-tagged wild-type or mutant DNMT3L constructs were harvested 56 days post-infection. Following trypsinization, cells were washed with PBS buffer and lysed in ice-cold M-PER™ mammalian protein extraction reagent (Thermo Scientific) supplemented with 2% Sodium dodecyl sulfate (SDS). The extracted proteins were denatured in SDS-loading buffer and analyzed by western blot using anti-Myc tag antibody (EMD Millipore 05724) and anti-β-tublin antibody (Cell Signaling Technology CST-86298).

### Illumina Infinium methylation EPIC array and data analysis

Illumina Infinium methylation EPIC array and data analysis were performed as previously described^10^. Briefly, genomic DNA was extracted from DKO8 cells 56 days post-infection with lentivirus carrying empty vector control, Myc-tagged wild-type or mutant DNMT3L constructs. DNA methylation was assessed using Illumina Infinium Human MethylationEPIC BeadChip array, conducted by the VAI Genomics core following the manufacturer’s specifications. Quality control, preprocessing and normalization process were performed using SeSAMe R package on Bioconductor^53^, and the methylation status of individual CpG site was reported as a β-value, ranging from 0 (unmethylated) to 1 (fully methylated). Changes of 0.3 (30%) β-value compared to empty vector were considered target sites for DNMT3L and its mutants.

