## Supplementary Figures for "Molecular Mechanisms of DNMT3A-3L-Mediated *de novo* DNA Methylation on Chromatin"

### Extended Data Fig. 1

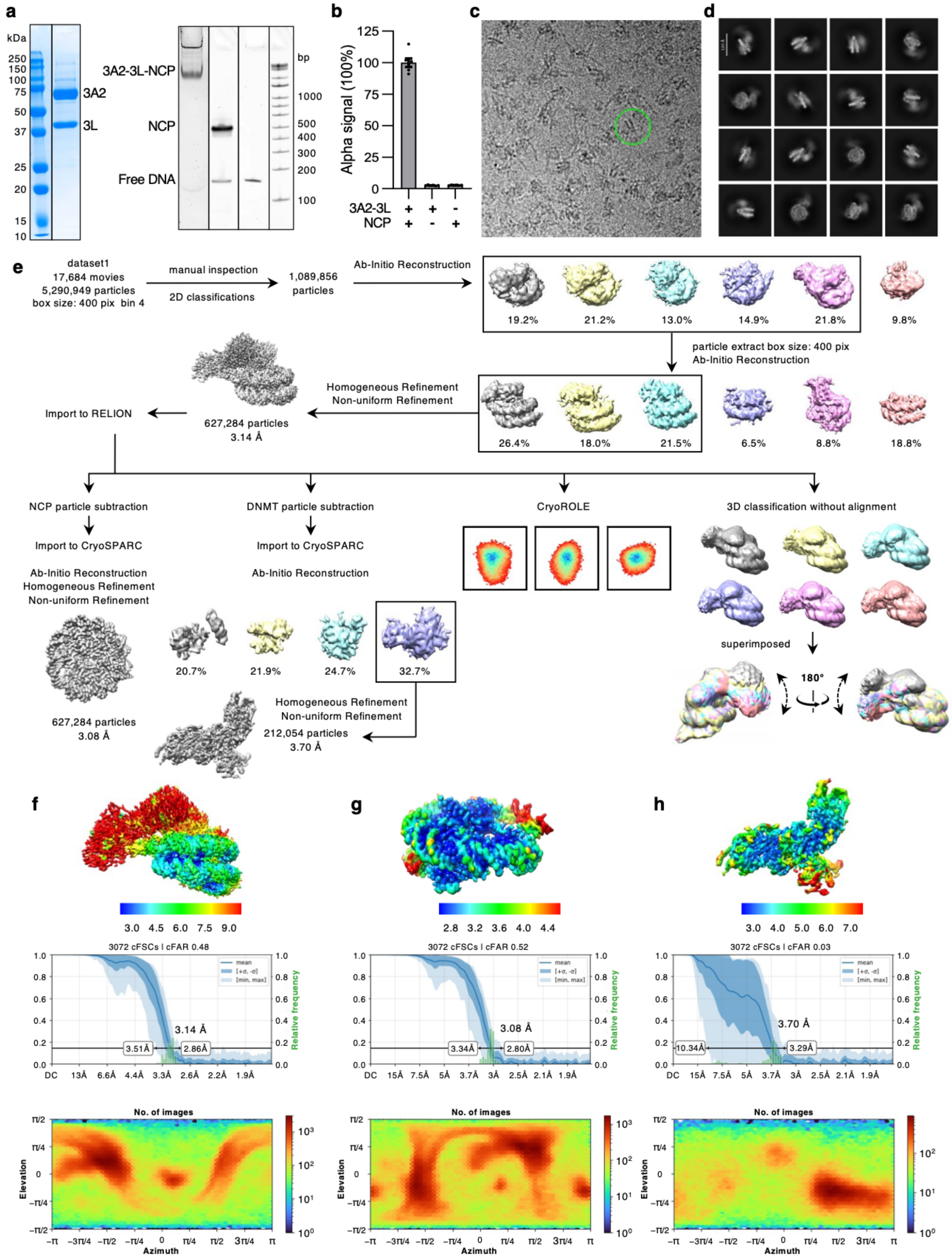

**Extended Data Fig. 1 Cryo-EM data collection and structure determination of DNMT3A2-3L-NCP complex from dataset1. a,** SDS-PAGE analysis of the DNMT3A2-3L complex and native gel analysis of the cross-linked DNMT3A2-3L-NCP complex. For gel source data, see **Supplementary Fig. 2**. **b,** AlphaScreen interaction assay between His-tagged DNMT3A2-3L and biotinylated NCPs. Data were presented as mean ± s.e.m., n = 6. **c,** Representative micrograph of DNMT3A2-3L-NCP complex with one particle highlighted in a green circle of 250 Å diameter. **d,** Representative 2D class-averages. **e,** Workflow diagram of cryo-EM data processing and 3D reconstructions in CryoSPARC (4.6.0) and particle subtraction in RELION 5. Boxed 3D classes were selected for further processing. The final global nominal resolution for the DNMT3A2-3L-NCP complex was 3.14 Å, for the NCP was 3.08 Å, and for the DNMT3A2-3L was 3.70 Å. Color-coded local resolution maps for consensus refinement (**f**), focused nucleosome refinement(**g**), and focused DNMT refinement(**h**). The final global nominal resolution was set at 3.14 Å for DNMT3A2-3L-NCP complex, 3.08 Å for NCP, and 3.70 Å for DNMT3A2-3L at 0.143 FSC. Half-map corrected Fourier Shell Correlation (cFSC) plots and angular distribution plots obtained from CryoSPARC (color indicative of the number of particles, increasing from blue to red, in a defined orientation) were shown at the bottom.

### Extended Data Fig. 2

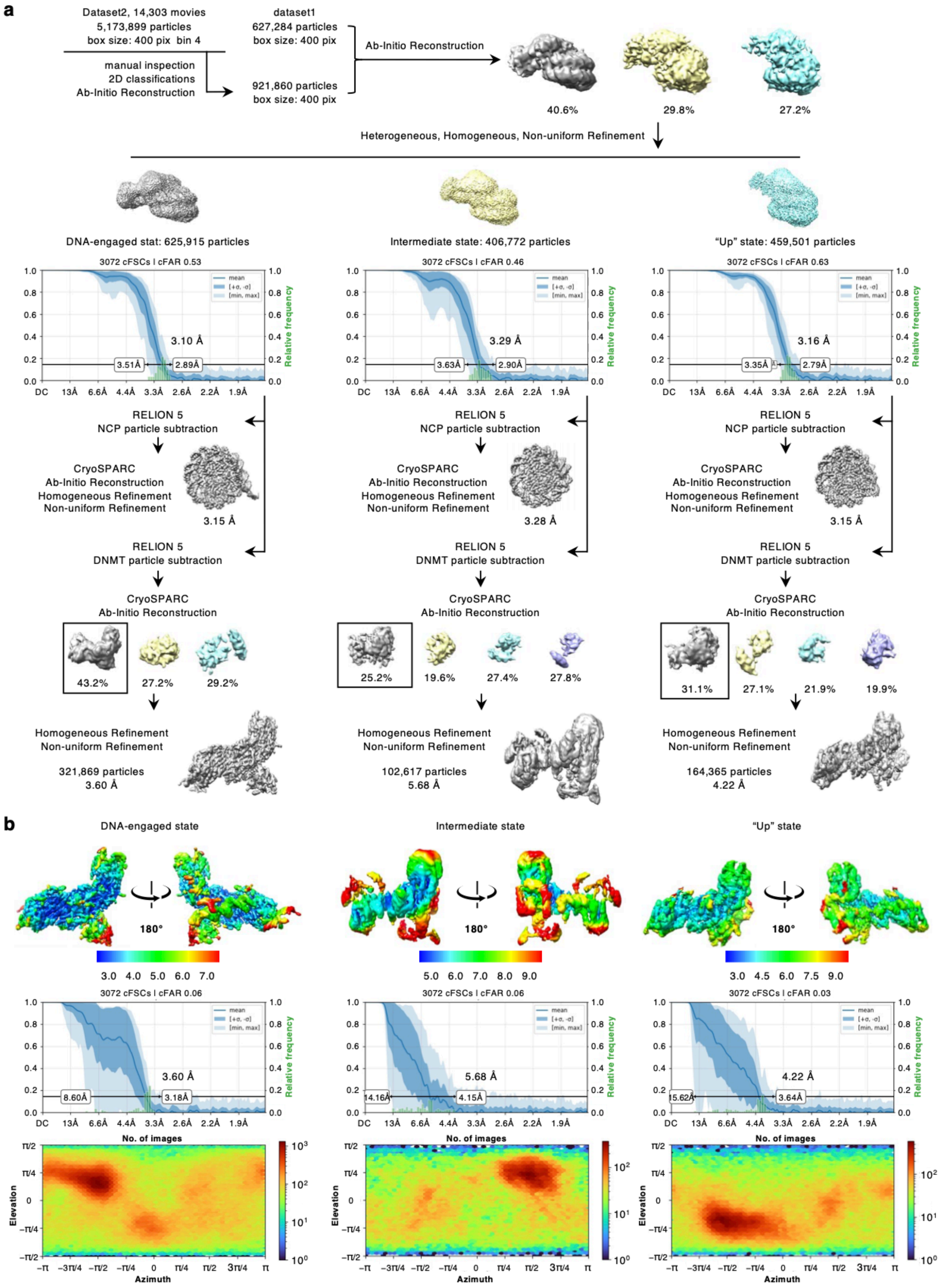

**Extended Data Fig. 2 Structure determination of DNMT3A2-3L-NCP in the combined dataset. a,** Cryo-EM data processing workflow by CryoSPARC (4.6.0) and RELION 5 of the combined dataset for the DNMT3A2-3L-NCP complex. Half-map cFSC plots were shown on the bottom for the three distinct conformations: a DNA-engaged state (grey, left), an intermediate state (yellow, middle), and an "up" state (cyan, right). The final global nominal resolution was set at 3.10 Å for the engaged state, 3.29 Å for the intermediate state, and 3.16 Å for the “up” state at 0.143 FSC. Particles from each conformation were further processed with focused refinements of DNMT and NCP separately. Boxed 3D classes were selected for further processing. The final global nominal resolution was set at 3.15 Å, 3.28 Å, and 3.15 Å at 0.143 FSC for NCPs from three conformations, and at 3.60 Å, 5.68 Å, and 4.22 Å at 0.143 FSC for DNMT3A2-3L from three conformations. **b,** Color-coded local resolution maps in two different orientations, related by a 180° rotation around a vertical axis for DNMT3A2-3L complexes in three distinct conformations. Half-map cFSC plots and angular distribution plots obtained from CryoSPARC were shown at the bottom for each conformation.

### Extended Data Fig. 3

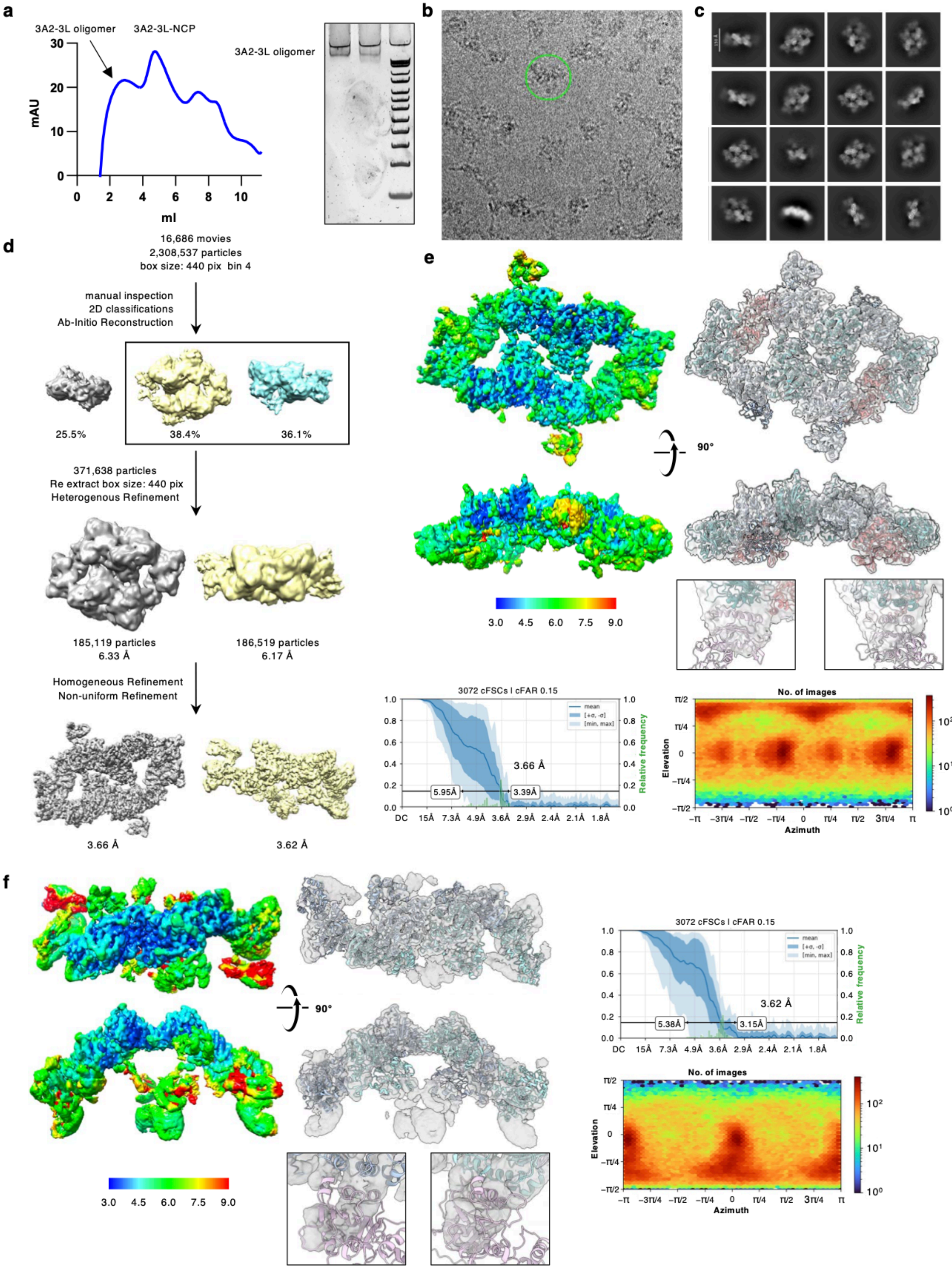

**Extended Data Fig. 3 Cryo-EM data collection and structure determination of DNMT3A2-3L oligomer complex. a,** The UV (280 nm) absorption profile of cross-linked DNMT3A2-3L-NCP sample collected from bottom to top (left). Native gel analysis of cross-linked DNMT3A2-3L oligomer complex (right). For gel source data, see **Supplementary Fig. 2**. **b,** Representative micrograph of DNMT3A2-3L complex with one particle highlighted in green circle of 300 Å diameter. **c,** Representative 2D class-averages. **d,** Workflow of cryo-EM data processing and 3D reconstructions in CryoSPARC (4.6.0). Boxed 3D classes were selected for further processing. The final global nominal resolution for the DNMT3A2-3L oligomer complexes was 3.66 Å for the dodecamer and 3.62 Å for the hexamer. **e,** Color-coded local resolution maps for DNMT3A2-3L dodecamer in two different orientations and overall atomic model fitting into the cryo-EM map density. Inlet, overlay of distal EM density with DNMT3L (pink) at lower contour, indicating the dodecamer. Half-map cFSC plots and angular distribution plots obtained from CryoSPARC were shown at the bottom. **f**, Color-coded local resolution maps for DNMT3A2-3L hexamer in two different orientations and overall atomic model fitting into the cryo-EM map density. Inlet, overlay of distal EM density with DNMT3L (pink) at lower contour, indicating further oligomerization. Half-map corrected cFSC plots and angular distribution plots obtained from CryoSPARC were shown on the right.

**Extended Data Fig. 4**

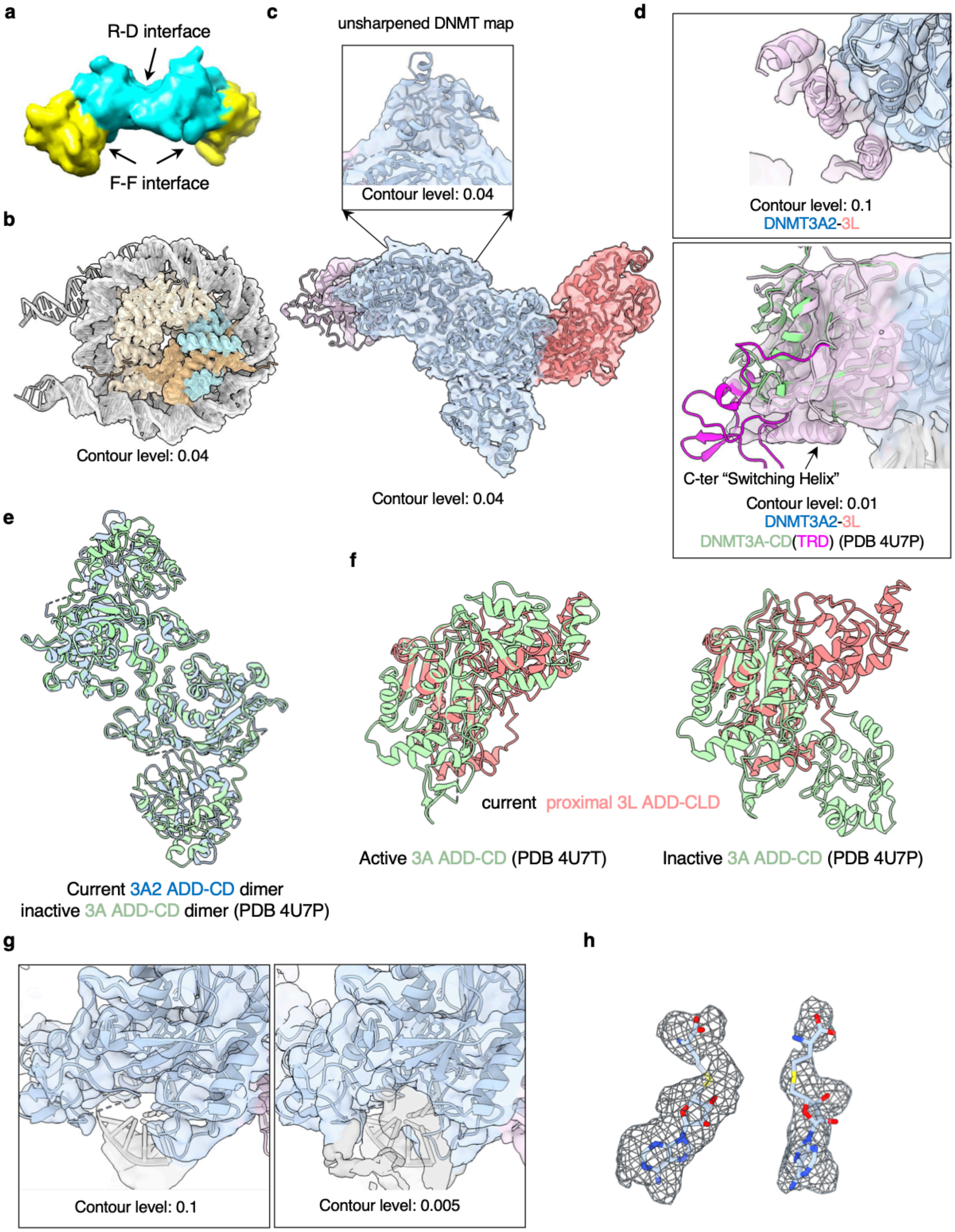


**Extended Data Fig. 4 Structure analysis of DNMT3A2-3L bound to nucleosomes. a,** Interfaces of the DNMT3A (CD)-3L(CLD) in crystal structure (PDB ID 2QRV). DNMT3A CD can form a V-shaped, polar, homodimeric interface (3A-3A RD interface) while with DNMT3L CLD, DNMT3A CD can form a flat, non-polar, heterodimeric interface (3A-3L FF interface) due to lack of the TRD loops in DNMT3L. **b,c,** Overall cryo-EM map fittings of NCP (**b**) and DNMT3A2-3L (**c**). Inlet, density map of the distal DNMT3A2 ADD domain generated in ChimeraX with the unsharpened map. **d,** Density map of the distal DNMT3L CLD domain generated in ChimeraX with contour level at 0.1 (top) and 0.01 (bottom). DNMT3L C-ter "Switching Helix" was indicated by the arrow. **e,** Structure comparison of DNMT3A2-3L-NCP with inactive DNMT3A ADD-CD dimer (PDB ID 4U7P). Only DNMT3A2 dimers were shown. **f,** Structure comparison of proximal DNMT3L with active (left) and inactive (right) DNMT3A ADD-CD domains (PDB ID 4U7T and 4U7P). **g,** Density map of the CD–DNA interaction region generated in ChimeraX with contour level at 0.1 (left) and 0.005 (right). **h,** Density maps of the two SAH ligands generated in ChimeraX and level at 0.05.

**Extended Data Fig. 5**

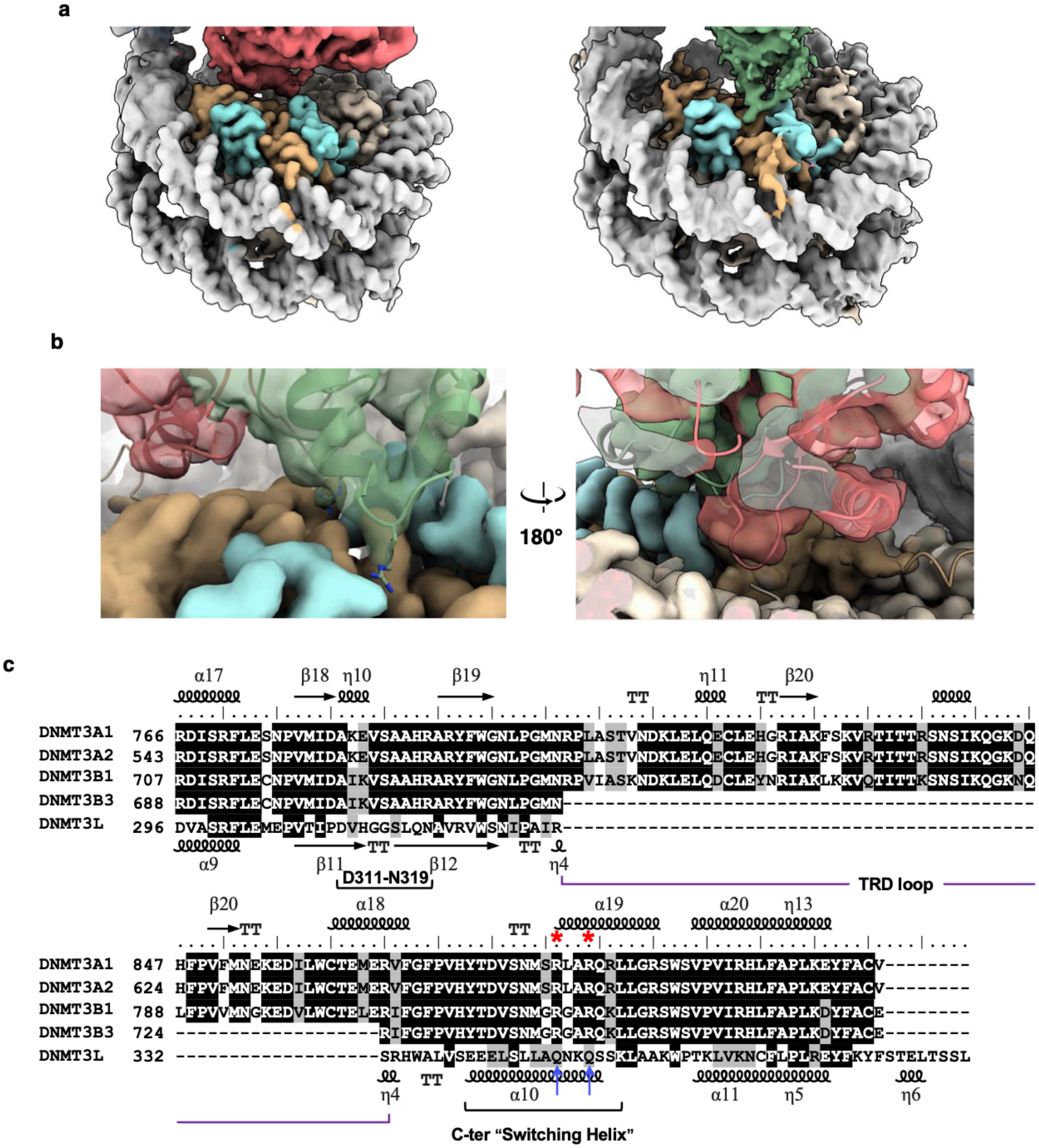


**Extended Data Fig. 5 Detailed structural insights into the interactions of DNMT3A2-3L with nucleosomes. a,** Cryo-EM densities of the nucleosome acidic patch region for the DNMT3A2-3L-NCP complex (left), and the DNMT3A2-3B3-NCP complex (right, EMD-20281). **b,** Two detailed views of superimposed maps of cryo-EM densities for the CLDs of DNMT3L and DNMT3B3 in the acidic patch region. The CLD regions were shown as cartoon models within the transparent density maps. **c,** Sequence alignments of the DNMT3 C-terminal region. Similar residues were shaded grey, while identical residues were shaded black. The TRD loop was missing in accessory proteins DNMT3L and DNMT3B3. The acidic patch-interacting arginine finger residues in DNMT3B3 (R740 and R743) were labeled with red stars, and the corresponding residues in DNMT3L (Q348 and Q351) were labeled with blue arrows. The alignment was annotated with secondary structure arrangements for DNMT3A1 (top) and DNMT3L (bottom).

**Extended Data Fig. 6**

**a**

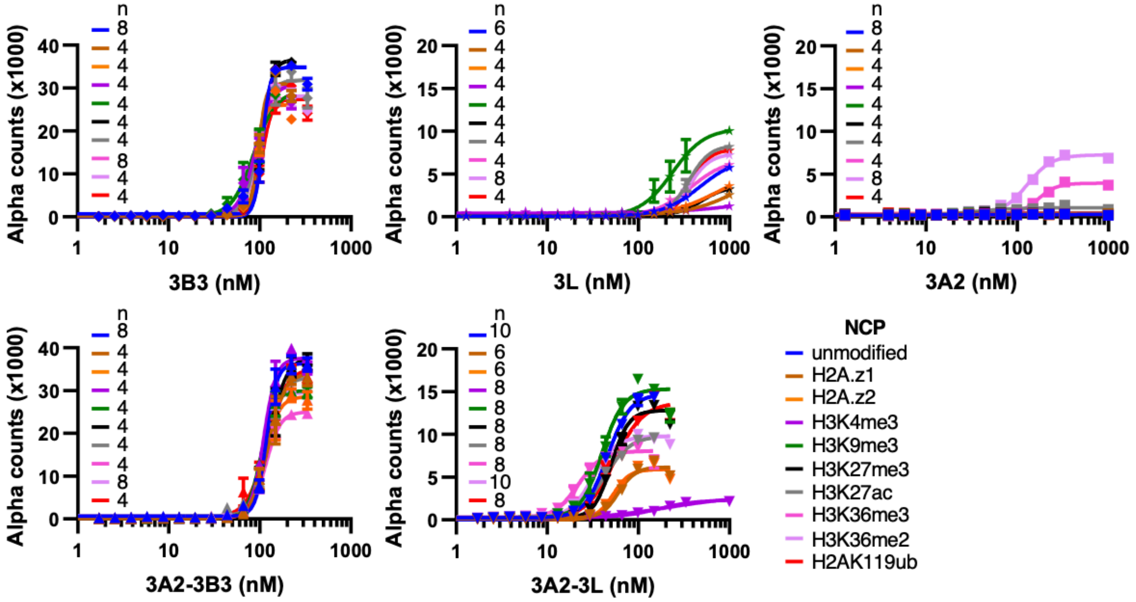


**b**

**$EC_{50}$ (nM)**

| NCP | 3A2 | 3B3 | 3L | 3A2/3B3 | 3A2/3L |
|---|---|---|---|---|---|
| unmodified | n.b. | 107 ± 5 | >472* ± 47 | 118 ± 6 | 46 ± 3 |
| H2AZ.1 | n.b. | 97 ± 3 | n.b. | 124 ± 11 | 61 ± 4 |
| H2AZ.2 | n.b. | 95 ± 5 | n.b. | 114 ± 9 | 54 ± 3 |
| H3K4me3 | n.b. | 98 ± 9 | n.b. | 110 ± 6 | 198 * ± 23 |
| H3K9me3 | n.b. | 86 ± 12 | 237 ± 68 | 103 ± 4 | 40 ± 4 |
| H3K27me3 | n.b. | 107 ± 3 | n.b. | 127 ± 11 | 50 ± 3 |
| H3K27ac | n.b. | 93 ± 5 | 359* ± 27 | 120 ± 6 | 42 ± 3 |
| H3K36me2 | 129 ± 9 | 94 ± 5 | 367* ± 25 | 114 ± 4 | 32 ± 2 |
| H3K36me3 | 170 ± 10 | 95 ± 6 | 375 ± 80 | 116 ± 4 | 22 ± 2 |
| H2AK119ub | n.b. | 106 ± 5 | >383 ± 33 | 119 ± 10 | 55 ± 5 |

**Binding**

Strong

Weak

No binding

n.b. : no binding; * : >1$\mu$M in some experiment repeats

**Extended Data Fig. 6 The interactions between DNMT3 complexes and nucleosomes. a,** AlphaScreen photo accounts for the interaction of his-tagged DNMT3A2, DNMT3A2-3B3, DNMT3A2-3L, DNMT3B3, and DNMT3L titrated against modified nucleosomes. Data were presented as mean ± s.e.m. n = 4-12 specified in each experiment. **b,** The full panel of DNMT3A2 nucleosomes binding strength ($EC_{50}$) in AlphaScreen assay.

**Extended Data Fig. 7**

**a**

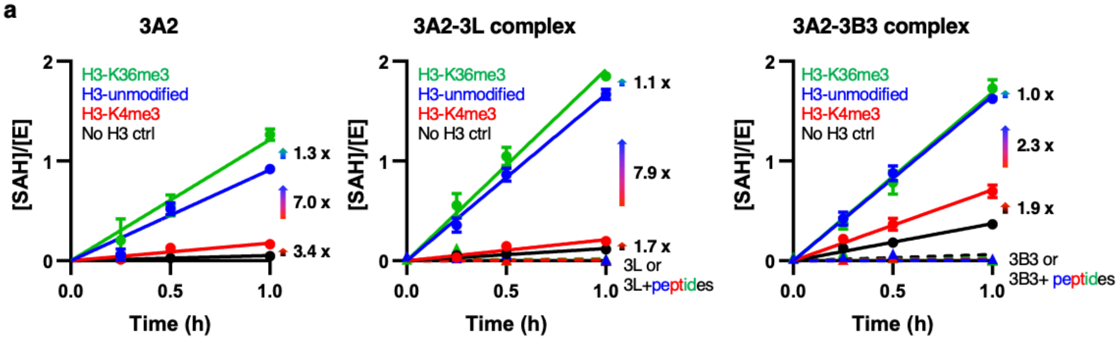


**b**

**Activities ([SAH]/[E]/h)**

| Enzyme | Histone H3 Peptides: unmodified | K36me3 | K4me3 | ctrl |
|---|---|---|---|---|
| 3A2 | 0.91 | 1.21 | 0.18 | 0.05 |
| 3A2-3L | 1.67 | 1.91 | 0.21 | 0.12 |
| 3A2-3B3 | 1.65 | 1.69 | 0.71 | 0.37 |
| 3L | -0.01 | 0.02 | 0.01 | 0.01 |
| 3B3 | 0.00 | 0.03 | 0.01 | 0.01 |

Activity: high – low

**d**

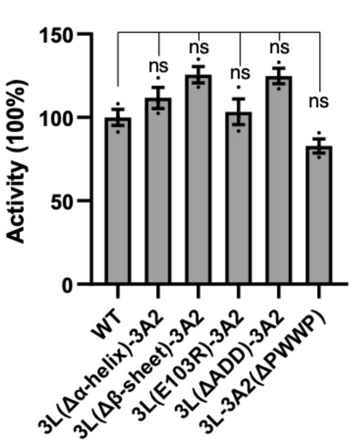


**c**

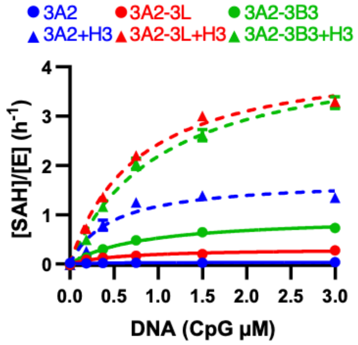


| DNMT3 | H3 peptide | Kcat (h-1) | Km (μM) | Kcat/Km (μM-1h-1) |
|---|---|---|---|---|
| 3A2 | - | 0.03 | 0.37 | 0.09 |
| 3A2-3L | - | 0.32 | 0.63 | 0.50 |
| 3A2-3B3 | - | 0.96 | 0.83 | 1.16 |
| 3A2 | + | 1.71 | 0.47 | 3.69 |
| 3A2-3L | + | 4.28 | 0.76 | 5.63 |
| 3A2-3B3 | + | 4.44 | 1.029 | 4.31 |

high – low

**e**

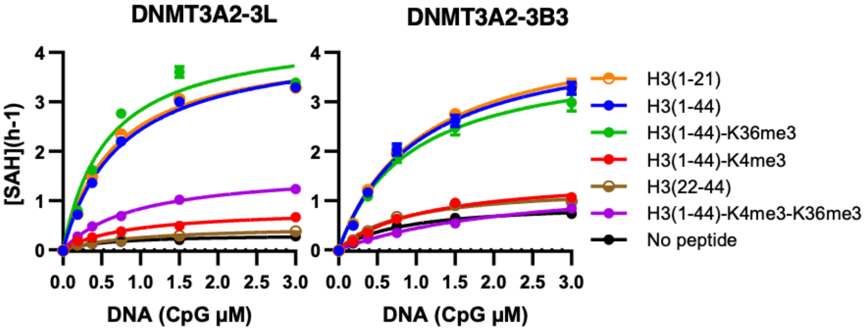


| DNMT3A2-3L Kcat (h-1) | DNMT3A2-3L Km (μM) | DNMT3A2-3L Kcat/Km (μM-1h-1) | DNMT3A2-3B3 Kcat (h-1) | DNMT3A2-3B3 Km (μM) | DNMT3A2-3B3 Kcat/Km (μM-1h-1) | Histone H3 peptide |
|---|---|---|---|---|---|---|
| 4.28 | 0.76 | 5.63 | 4.44 | 1.03 | 4.31 | H3(1-44) |
| 0.83 | 0.89 | 0.93 | 1.52 | 1.11 | 1.37 | H3(1-44)-K4me3 |
| 4.44 | 0.57 | 7.79 | 3.99 | 0.93 | 4.27 | H3(1-44)-K36me3 |
| 1.63 | 0.96 | 1.69 | 1.48 | 2.40 | 0.62 | H3(1-44)-K4me3-K36me3 |
| 4.19 | 0.67 | 6.29 | 4.60 | 1.06 | 4.33 | H3(1-21) |
| 0.49 | 0.99 | 0.50 | 1.30 | 0.81 | 1.61 | H3(22-44) |
| 0.32 | 0.63 | 0.50 | 0.96 | 0.83 | 1.16 | No peptide |

high – low

**Extended Data Fig. 7** ***In-vitro*** **DNMT3 DNA activities and kinetics**. **a,** *In-vitro* DNA methylation activities of DNMT3A2 (left), DNMT3A2-3L (middle), DNMT3A2-3B3 (right) in the absence (black) and presence of histone H3 peptide (1-44) unmodified (blue) or carrying K4me3 (red) or K36me3 (green). **b,** The full panel of DNMT3A2 DNA methylation activities with/without accessory proteins in the absence and presence of histone H3 peptide (1-44). **c,** Kinetics of DNMT3A2 with/without accessory proteins (DNMT3L or DNMT3B3) in the absence and presence of histone H3 peptide (1-44). **d**, *In-vitro* DNA methylation activity of wild-type or mutant DNMT3A2-3L complexes. Shown was the percentage methylation. **e,** Kinetics of DNMT3A2-3L and DNMT3A2-3B3 in the absence or presence of different H3 peptides. Data were presented as mean ± s.e.m., n = 8 (**a**) and 3 (**c, d, e**). ns, not significant. P-values, two-tailed Student's *t*-test.

**Extended Data Fig. 8**

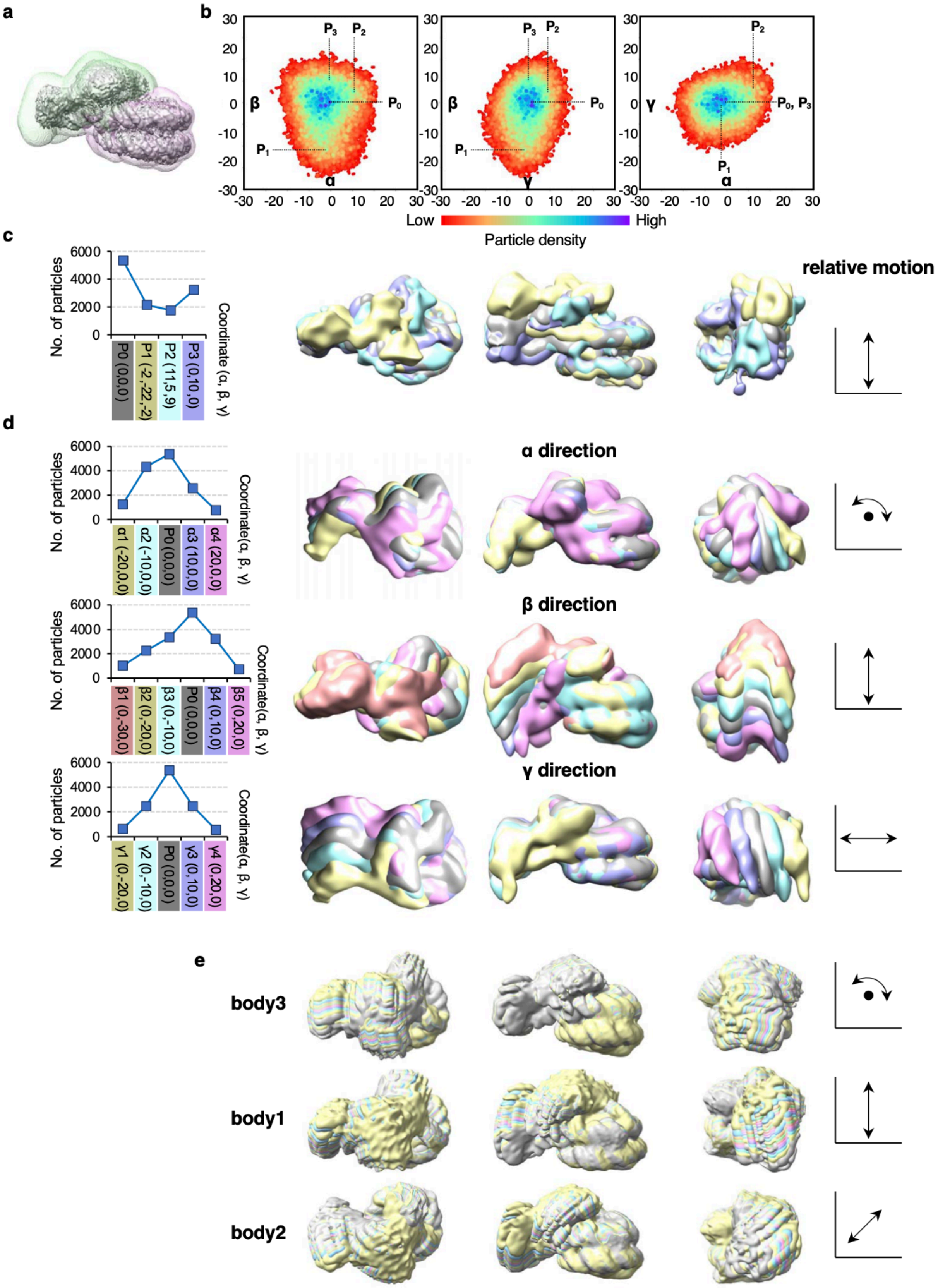

**Extended Data Fig. 8 CryoROLE and RELION5 multi-body analysis of the nucleosome-bound DNMT3A2-3L complex. a,** Particle signal subtraction masks for DNMT (green) and NCP (pink). **b,** 2D projection views of the conformational landscape generated using CryoROLE. **c,** Number of particles from 4 selected coordinates in the landscape (left) and superimposed views of four low-pass filtered (15 Å) 3D reconstructions calculated from particles within a 5 Å radius around these coordinates (middle). Relative motion trajectories were shown on the right. **d,** Overlays of 3D reconstructions calculated from particles selected along different directions (α, β, and γ) in the conformational landscape. The number of particles used was shown on the left. Relative motion trajectories were shown on the right. The reconstructions were calculated from locations at every 10° from +20° to -20° in α and γ direction, from +20° to -30° in β direction. All reconstructions were calculated from particles within a 5 Å radius around selected coordinates and low-pass filtered by 15 Å. **e,** Multi-body refinement in RELION5 showed similar motion between the nucleosomes and DNMT3A2-3L, especially the distal part of DNMT3A2-3L.

**Extended Data Table. 1** Cryo-EM data collection, refinement and validation statistics.

**Supplementary Fig. 1:** The gel source data for **Fig. 4e** and **Fig. 5d**. The boxed areas are the parts shown in the main figures.

**Supplementary Fig. 2:** The gel source data for Extended Data Figs. 1 and 3. The boxed areas are the parts shown in Extended Data Figs.

**Supplementary Fig. 3:** Calibration of SAH concentrations for MTase-Glo™ assay.

### Extended Data Table. 1

#### Cryo-EM data collection, refinement and validation statistics

| | DNMT3A2-3L-NCP data1 | DNMT3A2-3L-NCP data1+data2 consensus map (EMD-48322) | DNMT3A2-3L (EMD-48492) | NCP (EMD-48493) | DNMT3A2-3L-NCP intermediate state (EMD-48495) | DNMT3A2-3L-NCP "up" state (EMD-48494) | DNMT3A2-3L-NCP composed map (EMD-48498) (PDB 9MPP) | DNMT3A2-3L dodecamer (EMD-48496) (PDB 9MP0) | DNMT3A2 hexamer (EMD-48497) (PDB 9MPO) |
|---|---|---|---|---|---|---|---|---|---|
| **Data collection and processing** | | | | | | | | | |
| Microscope | | | | | FEI Titan Krios | | | | |
| Magnification | | | | | 105,000 | | | | |
| Voltage (kV) | | | | | 300 | | | | |
| Electron exposure (e–/Å$^2$) | | | | | 60 | | | | |
| Defocus range (μm) | | | | | -1.0 to -2.0 | | | | |
| Pixel size (Å) | | | | | 0.828 | | | | |
| Symmetry imposed | C1 | C1 | C1 | C1 | C1 | C1 | | C2 | C2 |
| Initial particle images (no.) | 5,290,949 | 10,464,848 | 625,915 | 627,284 | 10,464,848 | 10,464,848 | | 2,308,537 | 2,308,537 |
| Final particle images (no.) | 627,284 | 625,915 | 321,869 | 627,284 | 406,772 | 459,501 | | 185,119 | 186,519 |
| Map resolution (Å) | 3.14 | 3.10 | 3.60 | 3.08 | 3.29 | 3.16 | | 3.66 | 3.62 |
| FSC threshold | 0.143 | 0.143 | 0.143 | 0.143 | 0.143 | 0.143 | | 0.143 | 0.143 |
| Map resolution range (Å) | 3.51 ~ 2.86 | 3.51 ~ 2.89 | 8.60 ~ 3.18 | 3.34 ~ 2.80 | 3.63 ~ 2.90 | 3.35 ~ 2.79 | | 5.95 ~ 3.39 | 5.38 ~ 3.15 |
| **Refinement** | | | | | | | | | |
| Initial model used (PDB code) | | | | | | | 6PA7, 4U7P | 4U7P | 4U7P |
| Model resolution (Å) | | | | | | | 3.60 ~ 3.08 | 3.66 | 3.62 |
| FSC threshold | | | | | | | 0.5 | 0.5 | 0.5 |
| Model resolution range (Å) | | | | | | | 40 ~3.08 | 40 ~ 3.66 | 40 ~3.62 |
| Map sharpening *B* factor (Å$^2$) | -112.5 | -108.7 | -133.5 | -105.5 | -110.3 | -105.4 | | -120 | -125.7 |
| Model composition | | | | | | | | | |
| Non-hydrogen atoms | | | | | | | 23091 | 33333 | 18326 |
| Protein residues | | | | | | | 2023 | 4128 | 2264 |
| DNA base pairs | | | | | | | 332 | 0 | 0 |
| Ligands | | | | | | | 2 | 0 | 0 |
| Zn Ions | | | | | | | 6 | 30 | 12 |
| *B* factors (Å$^2$) | | | | | | | | | |
| Protein | | | | | | | 105.62 | 194.9 | 207.58 |
| Nucleotide | | | | | | | 132.4 | n/a | n/a |
| Zn | | | | | | | 191.87 | 260.95 | 298.13 |
| R.M.S. deviations | | | | | | | | | |
| Bond lengths (Å) | | | | | | | 0.008 | 0.006 | 0.007 |
| Bond angles (°) | | | | | | | 1.127 | 1.085 | 1.202 |
| Validation | | | | | | | | | |
| MolProbity score | | | | | | | 1.50 | 1.66 | 1.44 |
| Clashscore | | | | | | | 4.70 | 6.69 | 8.16 |
| Poor rotamers (%) | | | | | | | 0.06 | 0.47 | 0.3 |
| Ramachandran plot | | | | | | | | | |
| Favored (%) | | | | | | | 96.23 | 95.79 | 98.12 |
| Allowed (%) | | | | | | | 3.77 | 4.21 | 1.88 |
| Disallowed (%) | | | | | | | 0.00 | 0 | 0.00 |

**Supplementary Fig. 1: The gel source data for Fig. 4e and Fig. 5d.**

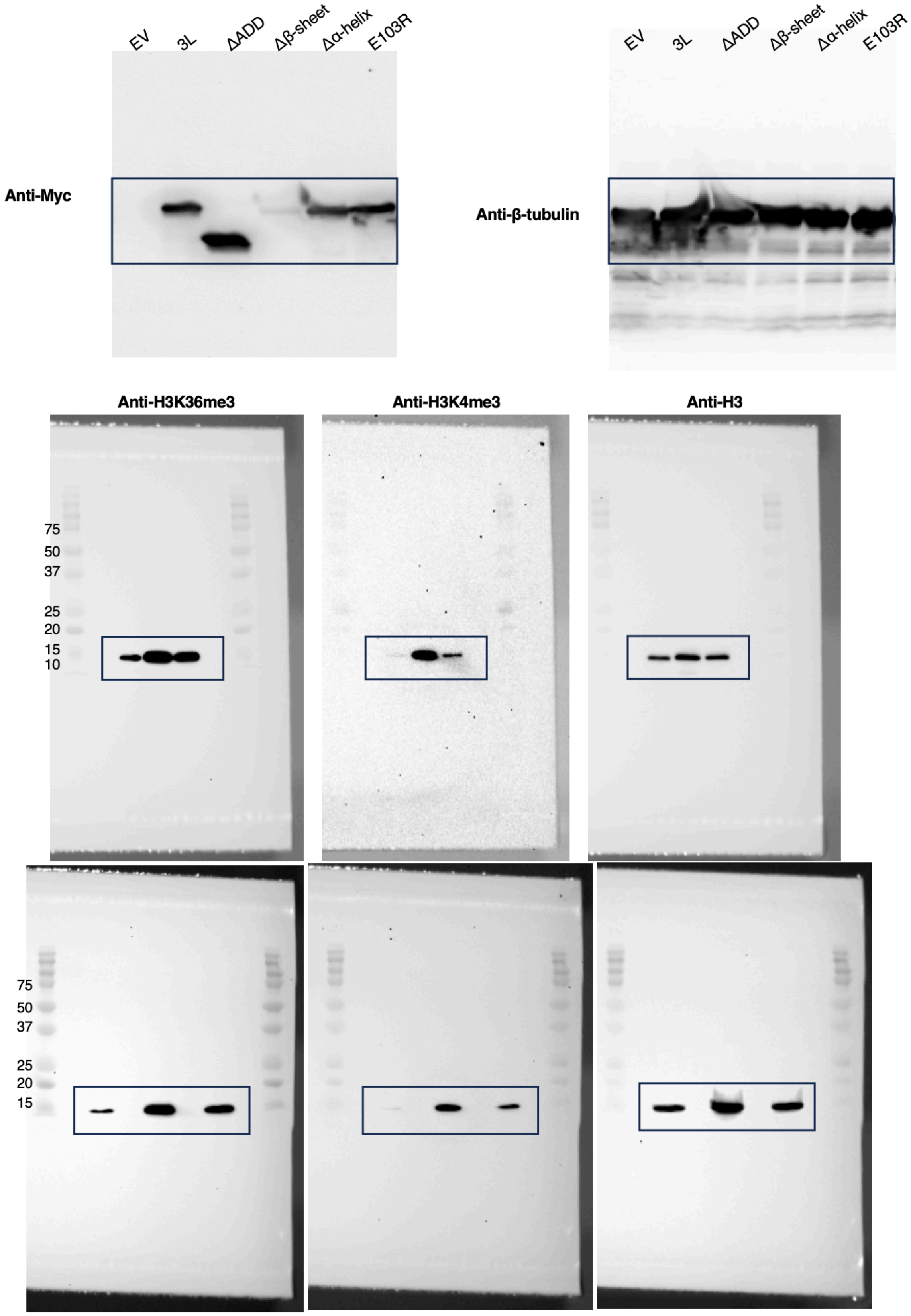

**Supplementary Fig. 2: The gel source data for Extended Data Fig. 1 and Extended Data Fig. 3.**

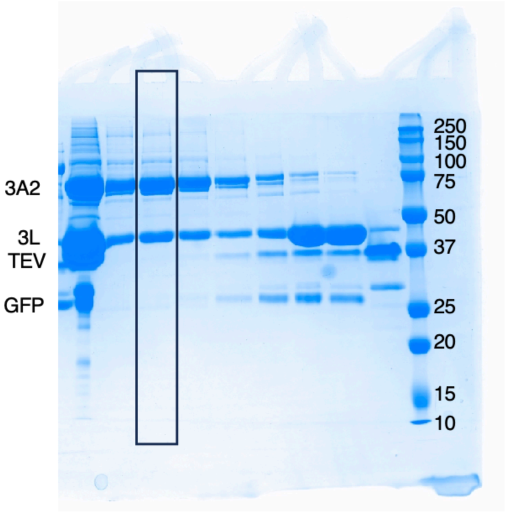


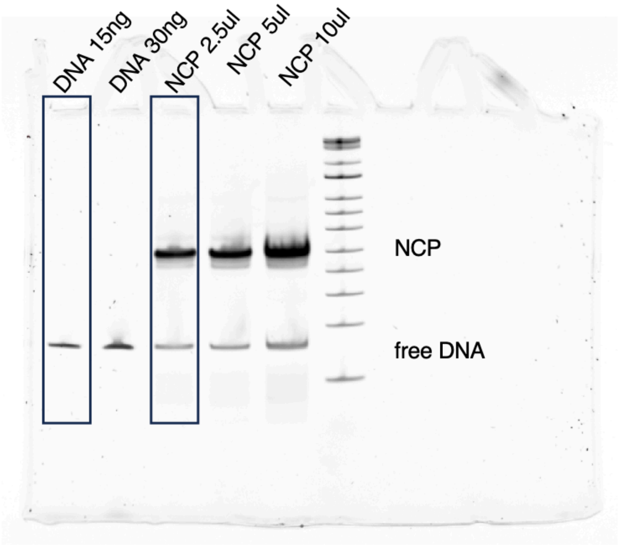


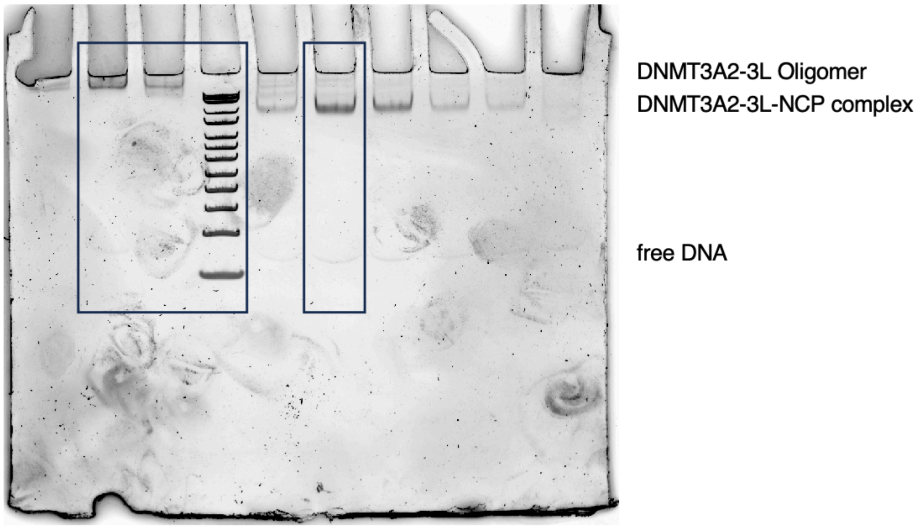

**Supplementary Fig. 3: Calibration of SAH concentrations for MTase-Glo™ assay.**

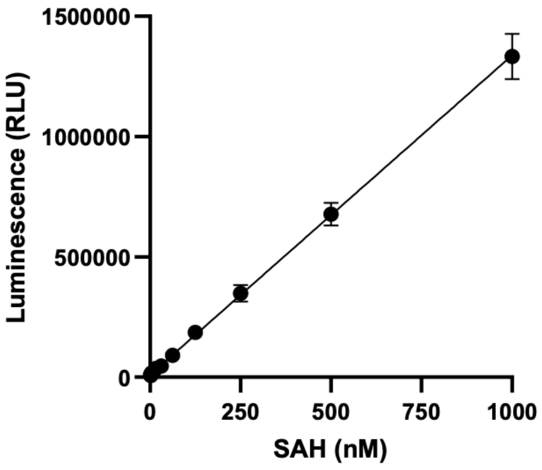


RLU = 1327.9077*[SAH] + 9679.2796
$R^2 = 0.9998$
